# Investigative power of Genomic Informational Field Theory (GIFT) relative to GWAS for genotype-phenotype mapping

**DOI:** 10.1101/2024.04.16.589524

**Authors:** Panagiota Kyratzi, Oswald Matika, Amey H Brassington, Connie E Clare, Juan Xu, David A Barrett, Richard D Emes, Alan L Archibald, Andras Paldi, Kevin D Sinclair, Jonathan Wattis, Cyril Rauch

**Affiliations:** School of Veterinary Medicine and Science, University of Nottingham, College Road, Sutton Bonington, LE12 5RD, UK; École Pratique des Hautes Études, PSL Research University, St-Antoine Research Center, Inserm U938, 34 rue Crozatier, 75012 Paris, France; Div. Genetics and Genomics, The Roslin Institute and Royal (Dick) School of Veterinary Studies, University of Edinburgh, Easter Bush, Midlothian EH25 9RG, Scotland, UK; Agriculture and Horticulture Development Board, Middlemarch Business Park Siskin Parkway, East Coventry CV3 4PE, UK; School of Biosciences, University of Nottingham, College Road, Sutton Bonington, LE12 5RD, UK; Shanghai Leadingtac Pharmaceutical Co., Ltd, 781 Cailun Road, China (Shanghai) Pilot Free Trade Zone, Pudong, Shanghai 201203, China; Centre for Analytical Bioscience, School of Pharmacy, University of Nottingham, Nottingham NG7 2RD, UK; Nottingham Trent University, 50 Shakespeare Street, Nottingham NG1 4FQ, UK; Centre for Mathematical Medicine and Biology, School of Mathematical Sciences, University of Nottingham, University Park, Nottingham NG7 2RD, UK

**Author notes:** Correspondence: Cyril Rauch.

**Keywords:** Complex traits, GIFT, genotype-phenotype mapping studies, GWAS

## Abstract

Identifying associations between phenotype and genotype is the fundamental basis of genetic analyses. Inspired by frequentist probability and the work of R.A. Fisher, genome-wide association studies (GWAS) extract information using averages and variances from genotype-phenotype datasets. Averages and variances are legitimated upon creating distribution density functions obtained through the grouping of data into categories. However, as data from within a given category cannot be differentiated, the investigative power of such methodologies is limited. Genomic Informational Field Theory (GIFT) is a method specifically designed to circumvent this issue. The way GIFT proceeds is opposite to that of GWAS. Whilst GWAS determines the extent to which genes are involved in phenotype formation (bottom-up approach), GIFT determines the degree to which the phenotype can select microstates (genes) for its subsistence (top-down approach). Doing so requires dealing with new genetic concepts, a.k.a. genetic paths, upon which significance levels for genotype-phenotype associations can be determined. By using different datasets obtained in *ovis aries* related to bone growth (Dataset-1) and to a series of linked metabolic and epigenetic pathways (Dataset-2), we demonstrate that removing the informational barrier linked to categories enhances the investigative and discriminative powers of GIFT, namely that GIFT extracts more information than GWAS. We conclude by suggesting that GIFT is an adequate tool to study how phenotypic plasticity and genetic assimilation are linked.

**NEW & NOTEWORTHY:** The genetic basis of complex traits remains challenging to investigate using classic GWASs. Given the success of gene editing technologies this point needs to be addressed urgently since there can only be useful editing technologies if precise genotype-phenotype mapping information is available initially. GIFT is a new mapping method designed to increase the investigative power of biological/medical datasets suggesting, in turn, the need to rethink the conceptual bases of quantitative genetics.

## INTRODUCTION

Identifying associations between phenotype and genotype is the fundamental basis of genetic analysis. The development of high-density genotyping and whole genome sequencing has enabled DNA variants to be directly identified and Genome-Wide Association Studies (GWASs) have become the method of choice for mapping genotype to phenotype in large populations of unrelated individuals. GWAS have been employed in many species, and especially in the study of human disease (1). By 2021 the NHGRI-EBI GWAS Catalog listed 316,782 associations identified in 5149 publications describing GWAS results (2). Additionally, extensive collection of data has been initiated through efforts such as the UK Biobank (3), Generation Scotland (4) and NIH *All of Us* research program (https://allofus.nih.gov/) in the expectation that large-scale GWAS will elucidate the basis of human health and disease and facilitate precision medicine.

While genomic technologies have advanced rapidly, statistical models used to analyze genetic data are still based on the models developed by Fisher more than 100 years ago (5, 6). GWASs essentially make use of the Fisher method of partitioning genotypic values by performing a linear regression of phenotype on marker allelic dosage (7). Regression coefficients estimate the average allele effect size, and the regression variance is the additive genetic variance due to the locus (8). However, an ongoing debate exists over whether the present analysis paradigm in quantitative genetics is at its limits for truly understanding complex traits, namely traits resulting from many genes each with very small effect size (9). As a result, one may wonder whether alternative statistical model(s) could be invented and used to determine genotype-phenotype mappings.

GWASs are fundamentally linked to frequentist probabilities that, defined through relative frequencies, determines the validity of statistical inferences. In practice, frequentist probabilities are generated through the grouping of data into bins or categories to generate a bar chart, that is then interpolated to create a distribution density function (DDF) in the continuum limit. The DDF is, in turn, used to determine statistical inferences including average, variance, p-value and so on. However, since the DDF approximates the bar chart (and not the converse), and that it is not possible to differentiate data from within any given group/category, the DDF is constructed mathematically on the implicit assumption that information is missing to differentiate data from within any given group/category.

The notion of ‘missing information’ can be legitimate and defined experimentally. For example, measuring the phenotype human height with a ruler with centimetre graduations implies that any height can be measured to the nearest centimetre. Consequently, one centimetre-width bins/categories need to be used to generate a frequency table of range of phenotype values upon which the phenotype and genotype DDFs are defined. In this case, all the resulting statistical inferences are defined with a precision corresponding to the nearest centimetre. The ‘missing information’ (i.e., that what cannot be measured by the ruler) corresponds then to sub-centimetric scales (i.e., distances to the nearest millimetre for this example). In practice the ‘missing information’ is therefore linked to the one of ‘imprecision’ and deciding to provide more precise statistical inferences implies that the width of categories be reduced, which can only be achieved by increasing the sample size. It is not by chance that the ‘normal distribution’ created by mathematicians and physicists was initially called the ‘law of errors’, where the notion of error (misinformation) results from imprecisions in experimental measurements. As a result, GWAS is faced with a fundamental issue involving the extraction of precise information using a method that, conceptually, assumes that information is missing or that data is mis-(in)formed.

In general, the problem concerning the ‘missing information’ is never mentioned since the DDF in the continuum limit is never considered as an approximation but as something that has its own reality. Namely a DDF must exist independently of data measured (i.e., data must fit the DDF and not the converse). The latter remark leads to an interesting conceptual territory where the notions of average and variance, and their usage, may be questioned. If one considers the normal distribution (or any other DDFs) is inherent to life and that data must fit it (them), then the moments of the distribution (e.g., average and variance) are also essential parameters to describe life, and the variance often interpreted as noise in the data is then a nuisance. If, on the contrary, data is the important thing, and that the DDF is considered solely as a tool to interpolate data based on missing information, then average and variance are parameters derived from a lack of information and are, as a result, poorly informative. The latter point should not come as a surprise as reducing the huge diversity of populations to a handful of parameters (i.e., average and variance) is highly reductionist and likely to be poorly descriptive. Thus, while the notions of average and variance may help representing datasets, they are inventions nonetheless, i.e., thought constructions akin to the field of frequentist probability. Thus, using average and variance as a starting point to map genotype-phenotype (GWAS) is a matter of choice. Accordingly, different statistical methods can be suggested.

To avoid those conceptual and practical issues a new method called GIFT (Genomic Informational Field Theory) has been designed and applied to simulated genotype-phenotype data in (10, 11), reviewed in (12). In short, to associate genotype to phenotype GIFT does not presume that the only important information concerning the gene effect is found in averages or variances, nor does it presume that DDFs are central. On the contrary, GIFT starts with the pre-requisite that phenotypic values, or phenotypic residuals after considering the environment/fixed effects, may be measured with sufficient precision to be unique in a population. Then, by avoiding grouping data into bins/categories, which would otherwise create an artificial imprecision, GIFT considers the entire information contained in the data, (i.e., variance is not a nuisance anymore) making use of the cumulative sum of microstates. Figure 1 provides the intuition underscoring GIFT as a method.

**Figure 1:**
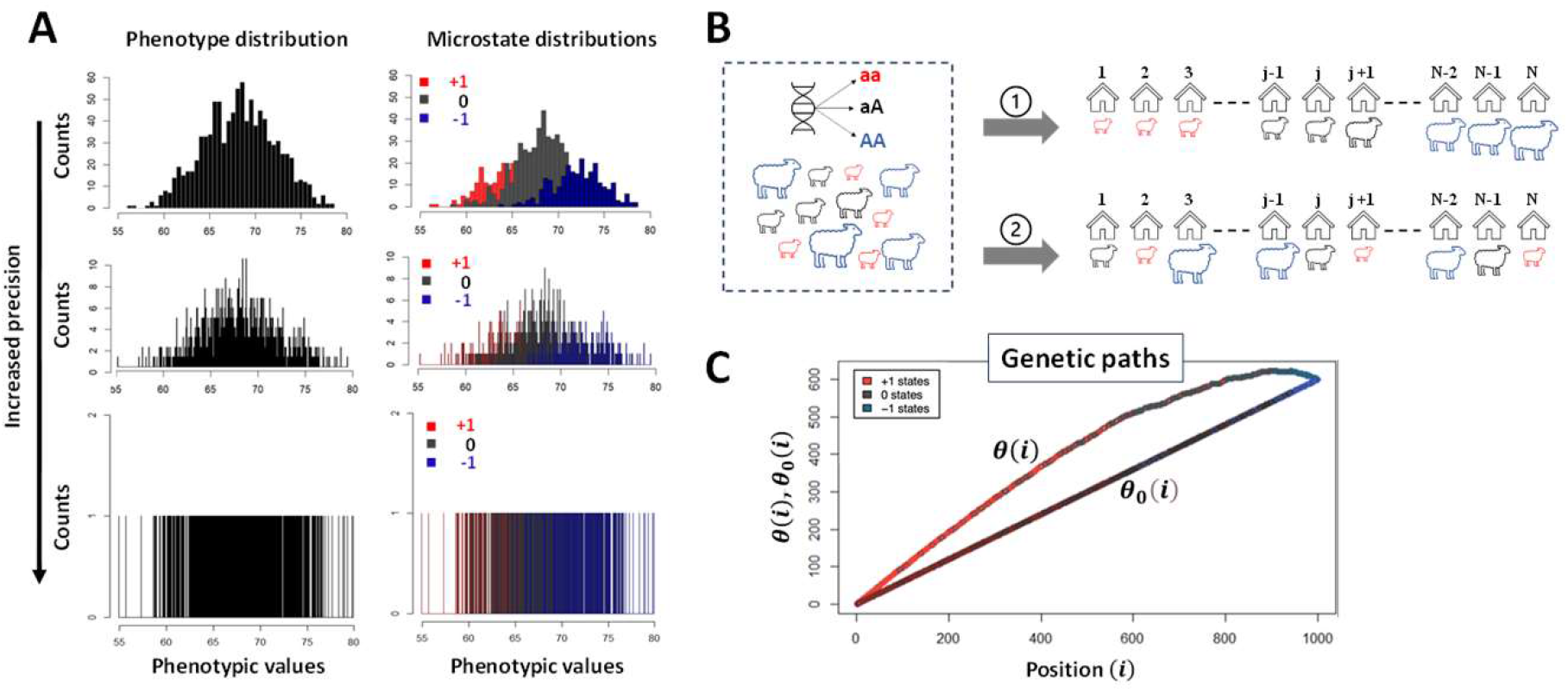
**(A)** For diploid organisms and for a binary (bi-allelic, A or a) genetic marker, any microstate (genotype) can only take three values that we shall write as ‘+1’, ‘0’ and ‘-1’ corresponding to genotypes aa, Aa and AA, respectively. The genotypes are color-coded to facilitate the representation of GIFT (+1: aa (red), 0: aA/Aa (black) and -1: AA (blue)). GWASs rely on probability density functions formed through the grouping of data into bins/categories. The phenotype distribution density function (A-top left) is then decomposed onto the distribution density function of genetic microstates (A-top right) for every single nucleotide polymophism (SNP). Using an analysis of averages and variances such decomposition determines whether the SNP studied is associated with the phenotype by comparing the average and variances of distributions. Repeating the same operation for every SNP in the genome permits to map genotype to phenotype. However, as more precise inferences can only come with, and are only legitimized by, a reduction in the width of categories, larger sample sizes are needed. To overcome this issue one way to proceed is to deconstruct density functions and wonder what would happen if one were able to reduce the width of categories, that is increasing the precision in the measurement of the phenotype or equivalently getting access to the whole information of datasets, without changing the sample sizes (**A** from top-to-bottom). The mathematical object that emerges is then a coloured barcode that is a list of microstates that can be analysed precisely by GIFT. **(B)** Such barcode can be obtained simply at the practical level through field studies. Assume a flock of sheep has been genotyped and that their phenotype has been measured sufficiently precisely such as to exclude the possibility that any two phenotypic values are identical. In the figure the magnitude of the phenotypic value for each sheep is characterised by the (unique) ‘size’ of the sheep. The barcode is obtained by ranking animals as a function of the magnitude of their phenotypic values (configuration ➀in Fig.1B). The null hypothesis is obtained via the random ranking of sheep that is equivalent to a lack of information on phenotypic values (configuration ➁ in Fig.1B). As GWAS works on phenotypic residual values after adjusting for fixed/environmental effects a similar barcode can be generated considering the magnitude of residual phenotypic values. **(C)** GIFT proceeds by plotting the cumulative sum of microstates as a function of the position in the list generating a curve called genetic path that is represented by θ(i) in Fig.1C and is unique to the SNP considered. While the curve θ(i) does not provide any significant information on its own, one may generate, for the same SNP, a curve (genetic path) corresponding to a sort of null hypothesis when ranking the phenotype does not bring any informational value. This is possible by scrambling (permutating) the string of microstates an infinite number of times. It is then possible to show that, in the asymptotic limit, the null hypothesis returns a straight line, noted θ_0_(i) (Fig.1C) out of which inferences may be suggested regarding potential association between the genotype and the phenotype by comparing θ_0_(i) to θ(i). Note, the simulation shown in **(A)** adhering to Fisher seminal model is based on a constant sample size of 1000 involving an arbitrary normally distributed phenotype of mean and variance 68 and 4 units, respectively. Each microstate is normally distributed with a gene effect identical to the standard deviation of the phenotype but without dominance. The frequency of the genotypes aa (red), Aa/aA (Grey) and AA (blue) are 64%, 32% and 4%, respectively and within Hardy-Weinberg ratio.

The current article extends our previous theoretic studies using simulated data to analyse for the first time two real datasets:

i. Dataset-1 is derived from a study concerned with the genetic background of carcass composition in sheep (*ovis aries*) (13). Using GWAS this study demonstrated a strong association between chromosome 6 and the carcass composition trait ‘bone area at the ischium’. We now apply GIFT to reanalyse this dataset to benchmark it against GWAS. Since GWAS previously identified a QTL in chromosome 6, our hypothesis was that GIFT would at least replicate GWAS results and identify additional putative QTLs.
ii. Dataset-2 comprises biochemical data arising from an ongoing study in sheep which seeks to identify risk allele variants in genes whose products direct a series of metabolic pathways, collectively referred to as one carbon (1C) metabolism and associated epigenetic regulators. The gene array was designed to include all single nucleotide polymorphisms (SNPs) linked to known biochemical enzymes involved in these pathways. Given that Dataset-2 preselected genes for a targeted analysis of enzymes involved in these metabolic/epigenetic pathways, it can be considered more specific.

The present article initially introduces the reader to the way data may be used and analysed differently using GIFT, contrasting to more conventional methods mostly based on an analysis of averages and variances. More specifically in Part 1, the null hypothesis defined by GIFT will be established. Using Dataset-1 the concept of genetic path pertaining to GIFT will be introduced (Part 2) out of which a p-value for GIFT will be defined (Part 3). Then Dataset-1 (Part 4) and Dataset-2 (Part 5) will be analysed comparing the informational/investigative power of GIFT relative to GWAS using Manhattan plots prior to performing enrichment analyses.

## MATERIALS AND METHODS

### Biological datasets

The first dataset (Dataset-1) analysed 600 pedigree-recorded Scottish Blackface lambs using CT scans to determine *in vivo* carcasses composition (13). The trait selected for the present study is the bone areas of the ischium (BAI) measured in mm^2^ from cross-sectional CT scans. The ischium is one of the three bones that make up the pelvis. It is located beneath the ilium and behind the pubis. The upper portion of the ischium forms a major part of the concave portion of the pelvis that forms the hip. The BAI crossed a genome-wide significance threshold on Chromosome 6 (OAR6). The pre-corrected phenotype values were obtained fitting fixed effects of age of dam, year of birth, the effect of management group (as sheep were from different farms), sex (males or females) and litter size (singles or twins) and as covariate the day of birth. Further information can be found in Matika et al. (2016) (13). Supplemental S1 provides the raw data used (Dataset-1).

The second dataset (Dataset-2) was from previously unpublished data extracted from a large ongoing programme of research to investigate genome regions (Quantitative trait loci, (QTL)) that determine metabolic and epigenetic responses to nutritionally induced deficiencies in one carbon metabolism (14, 15). For this study sheep were used as an experimental model. All animal procedures relating to this study adhered to the Animals (Scientific Procedures) Act, 1986. Associated protocols complied with the ARRIVE guidelines and were approved by the University of Nottingham Animal Welfare and Ethical Review Body (AWERB) with Home-Office project licensed authority (30/3376;10^th^ February 2016). Supplemental S2 provides the raw data used (Dataset-2).

### Dataset-2: Sheep genome resequencing, custom array design and SNP profiling on test subjects

Twenty-four unrelated Texel ewes were sequenced to a depth of 30x in 2 pools at Edinburgh Genomics. DNA samples were prepared using Illumina’s TruSeq PCR free kits and sequenced on an Illumina HiSeq 2500 Rapid Mode (serial no. D00125), read length of 150PE. Reads were trimmed to remove adapter sequences and low-quality bases using skewer with commands (-Q 20, -q 3) (16) and mapped to the reference sheep genome assembly (Oar_v3.1) using bwa mem (options -M -t 4) (17). Following deduplication using Picard-tools version 1.92, variants were called using GATK pipeline (18) including realignment around known indels and recalibration of bases, and FreeBayes (--use-best-n-alleles 4 -- pooled-discrete --min-alternate-count 4). Annotation of SNPs was performed using Ensembl variant effect predictor VEP version ensembl tools release 79 (19). 15,347,831 variants were identified. Of these, ∼3 million were novel SNPs and ∼12 million were already present in the Ensembl genome database. SNPs within annotated coding regions (VEP annotated “downstream gene variant” or “intron variant” removed) and within 3Kb upstream of a gene were retained. SNPs with a minor allele frequency of greater than 0.5 were used to design an Illumina Infinium® iSelect® Custom Array consisting of 4,576 probes. This captured SNPs in 115 1C metabolism and related genes, and 108 related epigenetic regulators as well as 33 control SNPs (Supplemental S1).

Liver samples were next collected post-mortem from 360 male and female Texel lambs (6 to 11 months of age) representing 11 farms dispersed regionally across the UK. Collections took place at regional abattoirs and samples immediately snap frozen in liquid N and stored at -80°C until analyses. DNA was then extracted using AllPrep DNA/RNA Mini kit (Qiagen, Manchester UK). Briefly approximately 20 mg of liver were mechanically disrupted using a TissueLyser (Qiagen, Manchester, UK) in 600 RLT plus buffer containing β-mercaptoethanol. Tissue lysates were then used to extract RNA and DNA according to the manufacturer instructions. The custom designed array was then used to SNP profile DNA from these Texel-sheep. For this purpose, liver samples were collected post-mortem from lambs (aged 6 to 11 months) representing 11 farms dispersed regionally across the UK. Collections took place at regional abattoirs and samples immediately snap frozen in liquid N and stored at -80°C until analyses. DNA was then extracted using AllPrep DNA/RNA Mini kit (Qiagen, Manchester UK). Briefly approximately 20 mg of liver were mechanically disrupted using a TissueLyser (Qiagen, Manchester, UK) in 600 RLT plus buffer containing β-mercaptoethanol. Tissue lysates were then used to extract RNA and DNA according to the manufacturer instructions.

### Dataset-2: Metabolic profiling

For the purposes of the current study the following seven liver metabolites were selected from a larger pool of 1C metabolites: S-adenosyl methionine (SAM), methylcobalamin (mB12), adenosylcobalamin (aB12), trimethylglycine (TMG), dimethylglycine (DMG), propionate (PPA) and methylmalonic acid (MMA). The first four metabolites were selected as representative intermediates of the methionine cycle whilst the latter two are intermediates in the hepatic synthesis of succinate (15) (Fig.2 & Supplemental S1).

**Figure 2:**
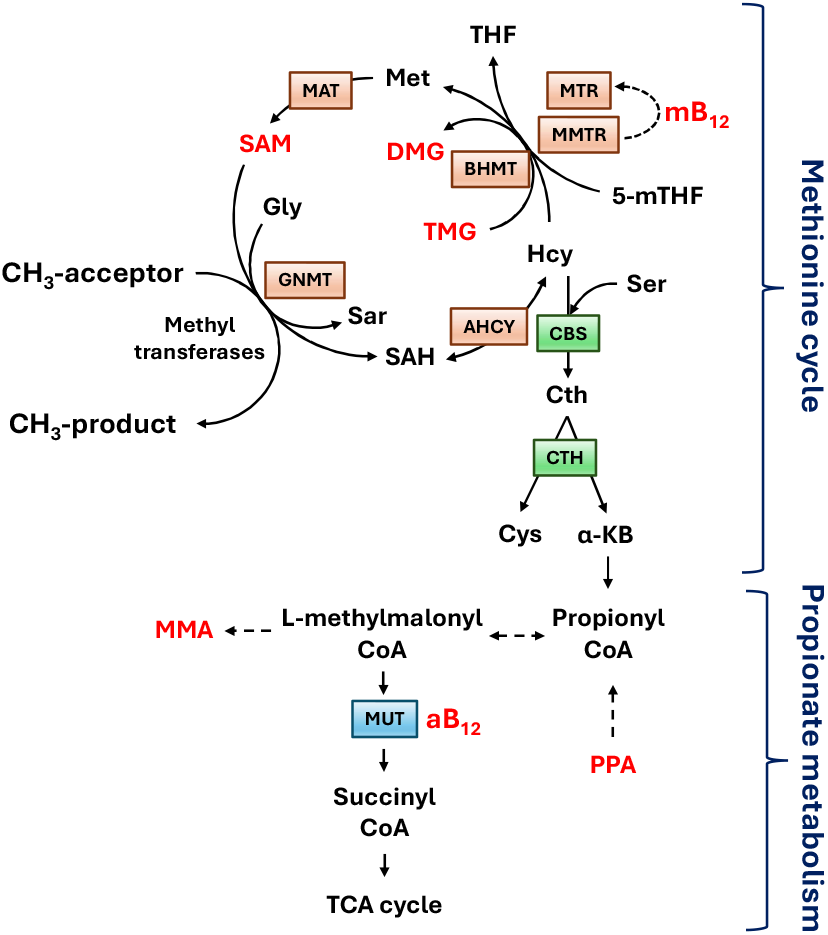
Linked methionine and propionate metabolism adapted from Clare et al. (2019) where all metabolites studied for this study are in red. The methionine cycle facilitates the re-methylation of homocysteine (Hcy) to methionine (Met) and ultimately S-adenosylmethionine (**SAM**) with methyl (CH_3_) groups donated either from folate (5-mTHF) or betaine (trimethylglycine; **TMG**), thus leading to the formation of dimethylglycine (**DMG**). Methylcobalamin (**mB12**) serves as a cofactor for the reduction of the inactive form of methionine synthase to its active state (MTR), which then transfers a methyl group from 5-mTHF to Hcy. The linked metabolism of propionate (**PPA**) to succinate (an intermediary metabolite in the tricarboxylic cycle) requires adenosylcobalamin (**aB12**), which serves as a cofactor for methylmalonyl-CoA-mutase (MUT) leading to the generation of succinyl-CoA and methylmalonic acid (**MMA**) in this pathway. Other intermediary metabolites and enzymes listed: glycine (Gly), sarcosine (Sar), S-adenosylhomocysteine (SAH), tetrahydrofolate (THF), serine (Ser), cystathionine (Cth), cysteine (Cys), alpha-ketobutyrate (α-KB), methylmalonic acid (MMA); Betaine homocysteine methyltransferase (BHMT), Methionine adenosyl-transferase (MAT), Glycine methyl-transferase (GNMT), Adenosyl-homocysteinase (AHCY), Cystathionine beta-synthase (CBS), cystathionine gamma-lyase (Cth).

Hepatic concentrations of four metabolites (i.e., mB12, aB12, TMG and DMG) were determined by hydrophilic interaction chromatography (HILIC) coupled to electrospray ionization tandem mass spectrometry (MS/MS) as reported previously (20). For the analysis of SAM (determined separately by HILIC), the standard was purchased from Sigma-Aldrich (Poole, Dorset, UK). Stock solutions of this standard were prepared in potassium phosphate extraction buffer (KH_2_PO_4_ and K_2_HPO_4_; 40 mmol/L) containing 0.1% L-ascorbic acid, 0.15% citric acid and 0.1% MCE (adjusted to pH 7 with NaOH), each at a final concentration of 100 μmol/L. Also, for SAM the mobile phase was modified from that used for the three other reported metabolites by adjusting the pH of the aqueous ammonium carbonate buffer solution from 3.5 to 9.1. Mass spectrometer parameters for SAM were as follows: retention time = 7.69 min; Q1mass = 399.1 amu; Q3 mass = 250.1 amu; declustering potential = 56; collision energy = 25; collision cell exit potential = 16.

Hepatic concentrations of PPA and MMA were determined by gas chromatography coupled to mass spectroscopic-detection (GC-MS). Briefly, for PPA, 750 μL 5-Sulfosalicylic acid (SSA, 0.04 mg/ml) was added to 150mg frozen liver, homogenised for 2 min and cooled on ice for 10 min. The sample was centrifuged for 15 min at 14,500 x g and 200 μL liver homogenate transferred to a 2.5 mL screw capped glass vial. To this, 20 μL internal standard (MBA, 400 μM), 3.5 μl HCl (37%) and 1 mL diethylether were added, vortexed for 2 min and centrifuged for 10 min at 14,500 x g. 600 μL of the upper layer was transferred to a screw capped glass vial containing 3.5 μL 1-(tert-butyldimethylsilyl)imidazole (TMDMSIM, 97%), vortexed for 2 min and heated at 60°C for 30 min. GC-MS analysis proceeded after cooling. The method used a DB-5MS column (J&W Scientific Agilent technology, 30 m x 0.25 mm; 0.25 μm film thickness). The carrier gas (He) was set at a constant flow rate of 1.3 ml/min. The injection volume was 5 μL for SCAN mode (for qualification) and SIM (selected ion monitoring) mode (for quantification), both using splitless mode. The injection port and MS selective detector interference temperatures were 260°C and 250°C respectively. The chromatograph was programmed for an initial temperature of 40°C for 1 min, increased to 60°C at 70°C min-1, then to 110°C at 15°C min-1, and finally 250°C at 70°C min^-1^. MS was tuned regularly and operated in electron impact (EI) ionization mode with the ionization energy of 70eV. SCAN mode measured at m/z: 30-300 and SIM ions were set at 159 (for MBA) and 131 (for PPA). The same method was used to produce a calibration curve for PPA using standards at concentrations ranging from 19.5 nmol/g to 5μmol/g. The limit of detection was 19.5 nmol/g. CVs for low, medium and high QCs were 10.4, 6.3 and 6.5% and the inter-assay CV was 4.7%.

For MMA, 250 μL 80% MeOH was added to 50 mg frozen liver, homogenised for 2 min and cooled on ice for 10 min. The sample was ten centrifuged for 15 min at 14,500 x g and 200 μL liver homogenate transferred to a 2.5 mL screw capped glass vial. To this, 4 μL internal standard (1 mM 4-chlorobutyric acid (CBA) in 1 mM HCl) followed by 250 μL 12% BF3-Methanol were added, vortexed for 1 min and heated at 95°C for 15 min. After cooling, 250 μL cold distilled water and 250 μL cold dichloromethane (CH_2_Cl_2_) were added to the vial, vortexed for 30s and centrifuged for 10 min at 14,500 x g. The lower dichloromethane layer was transferred to a screw capped glass auto-sampler vial with insert for GC-MS analysis. The method used a DB-WAX column (cross-linked polyethylene glycol; J&W Scientific Agilent technology) (30 mm x 0.25 mm; 0.15 μm film thickness). The carrier gas (He) was set at a constant flow rate of 1.0 ml/min. The injection volume was 1 μL for SCAN mode (for qualification) and SIM mode (for quantification), both using splitless mode. The injection port and MS selective detector interference temperatures were 260°C and 280°C respectively. The chromatograph was programmed for an initial temperature of 50°C for 2 min, increasing to 150°C at 8°C min-1, then to 220°C at 100°C min^-1^ and held for 5 min at the final temperature. MS was tuned regularly and operated in EI ionization mode with the ionization energy of 70eV. The limit of detection was 0.75 nmol/g for both MMA and SA and inter-assay CVs were 8.4% for MMA and 11.0% for SA.

### Dataset-2: Determination of GWAS for 1C-metabolites

Preliminary data analysis indicated the need to log-transform using the natural logarithm (Supplemental S3) to approximate normality. Transformed data were then pre-corrected for the fixed effects of farm (F) and sex (S) in ASReml using the following model, y_ij_ = μ + F_i_ + S_j_ + e_ij_, where y_ij_ is the log-transformed phenotype, that is the log-transformed metabolite concentration studied; μ is the overall mean for the log-transformed metabolite concentration; F_i_ is the effect of the i^th^ farm (i =1,..,11); S_j_ the effect of j^th^ Sex (Male vs Female) and, e_ij_ is the residual. The genotype dataset was filtered using PLINK (HWE p-value threshold of 10^−6^, call rate for genotypes of 10% and a MAF of 5%), the number of independent SNPs was determined using BCFTOOLS (r^2^-threshold=0.1) and the GWAS Manhattan plots, linked to the determination of p_GWAS_, were obtained using GEMMA. The same genotype and residual phenotypes as filtered by GWAS were used by GIFT.

### Data representation using GIFT

Adjusted phenotypic data (i.e., residuals, from Dataset-1 and Dataset-2) were used for this study. Regarding the representation of GIFT, upon selecting a SNP for all individuals, the different corresponding genotypes, aa, aA/Aa and AA, were assigned the arbitrary values +1, 0 and -1, respectively. With this convention any barcode can be represented by a string of numbers from which a GIFT analysis can be inferred. More specifically, the assignment of values +1, 0 and -1 were done as a function of the base pairs as follow: AA=TT=+1, GG=CC=-1 and 0 otherwise. As shown schematically in Fig.1, the residuals obtained were ranked by order of magnitude and the cumulative sum of their corresponding genotypic values performed to obtain the ‘genetic path’ for the SNP considered. The genetic path of a SNP is noted *θ*(*i*) in the text (Fig.1). The null hypothesis for GIFT as well as the notion of significance when GIFT is used will be introduced and fully explained in the RESULTS section.

## RESULTS

### Analyze of the null hypothesis θ_0_(i) for GIFT

While θ(i) is obtained using phenotypic information (configuration ➀ in Fig.1 and ‘Data representation using GIFT’ in MATERIALS AND METHODS), it is also possible to plot the cumulative sum of microstates when no phenotypic information is present that is equivalent to ‘scrambling’ or permutating the string of microstates in Fig.1A also corresponding to the configuration ➁ in Fig.1B. Recall that since our focus is on a given SNP, then the number of microstates, N_+_, N_0_ and N_−_, are identical between the configurations ➀ and ➁. This new cumulative sum noted θ_0_(i) is expected to be a sort of null hypothesis solely dependent on the bulk microstate frequencies N_+_/N, N_0_/N and N_−_/N, where N_q_ q ∈ {+,0, −} is the number of microstates of type q. This is so because there is no further information that could inform on the positioning of microstates in their list when the scrambled state is considered. However, while θ(i) is unique since phenotypic information is used to generate it, θ_0_(i) is not as each time the string of microstates from Fig.1A is scrambled, a new θ_0_(i) appears. Accordingly, one needs to consider the set of possible θ_0_(i)s generated bounded to the microstate frequencies N_+_/N, N_0_/N and N_−_/N.

Using a selection of theoretic SNPs defined by different microstate frequencies (Table-1). Fig.3A illustrates the global shape resulting from simulating 1000 θ_0_(i)s. The results demonstrate that the global shape of the θ_0_(i)s plotted as a function of the position in the string is ellipsoidal with short and long axes changing as a function of microstate frequencies involved, and where the different averages of θ_0_(i)s represented by black lines in Fig.3A, are straight lines with slopes linked to the difference, ΔN/N = (N_+_ − N)/N. The fact that the averages of θ_0_(i)s for a given set of microstates, N_+_, N_0_ and N_−_, is always a straight line linked to microstate frequencies, N_+_/N, N_0_/N and N_−_/N, can be understood intuitively by the fact that scrambling or permutating an infinite number of times the string of microstates is equivalent to determining, for any position i, the presence probability, N_q_/N, of each microstate in the string. Accordingly, for a given set of microstates, N_+_, N_0_ and N_−_, the average of θ_0_(i)s, noted ⟨θ_0_(i)⟩, is 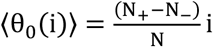. Further theoretic details can be found in (10, 11). Using ⟨θ_0_(i)⟩ as a reference for the null hypothesis, Fig.3B show the sur-imposition of the differences, Δθ_0_(i) = θ_0_(i) − ⟨θ_0_(i)⟩, obtained from simulations using SNPs from Table-1.

**Table 1:**
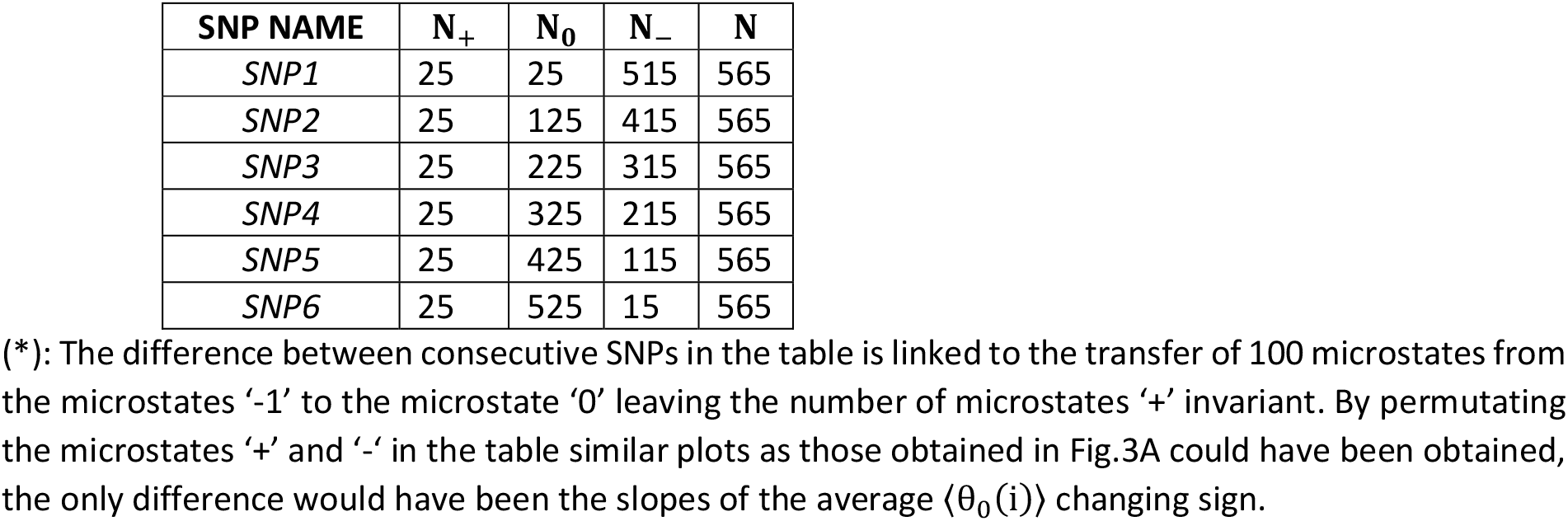
Theoretics SNPs used to capture the null hypothesis associated with GIFT upon 1000 simulations of microstates permutation*.

| SNP NAME | $N_+$ | $N_0$ | $N_-$ | N |
| --- | --- | --- | --- | --- |
| *SNP1* | 25 | 25 | 515 | 565 |
| *SNP2* | 25 | 125 | 415 | 565 |
| *SNP3* | 25 | 225 | 315 | 565 |
| *SNP4* | 25 | 325 | 215 | 565 |
| *SNP5* | 25 | 425 | 115 | 565 |
| *SNP6* | 25 | 525 | 15 | 565 |

**Figure 3:**
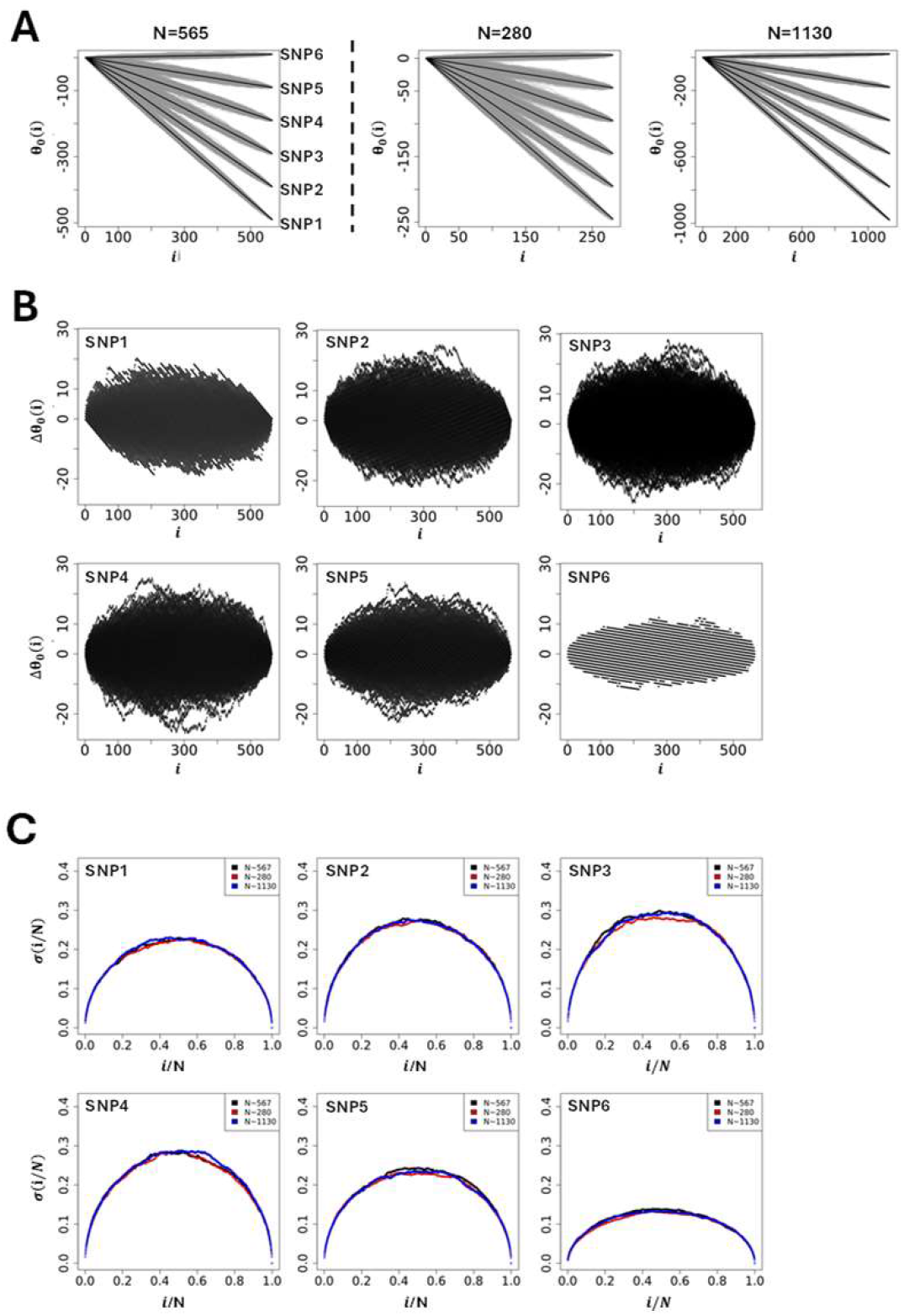
**(A-left panel)** Simulations of genetic paths corresponding to null hypotheses using GIFT as a method. The data used for the simulation are given in Table-1. **(A-right panel)** Simulations of genetic paths corresponding to null hypotheses when the sample size is divided or multiplied by a factor two. **(B)** Representation of Δθ_0_(i) = θ_0_(i) − ⟨θ_0_(i)⟩ for the microstates data as given in Table-1. **(C)** Plots of the standard deviation normalised by the square root of the sample size and where the position is also normalised by the sample size. The code for the simulations is given in Supplemental S4.

Finally, to assess the impact of the sample size (population size) on the null hypothesis the initial size (N=565, Table-1) was divided (N=280) and multiplied (N=1130) by a factor ∼2 while keeping constant the microstate frequencies N_+_/N, N_0_/N and N_−_/N from Table-1. The simulations Fig.3A show that the appearance of ellipsoids is affected when the sample size changes, becoming thinner as the population size increases. Plotting the standard deviation, σ(i/N), as a function of the position once normalized by the sample size, 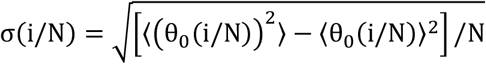, resulting from the different simulations in Fig.3C demonstrates that the standard deviation from GIFT is quadratic, and independent of the sample size, as expected from a random allocation of different microstates in the string of positions.

At first sight and with this primary analysis one could suggest that any genetic path departing from the cloud of genetic paths formed by the set of θ_0_(i)s upon the permutation of microstates (grey surface in Fig.3A or black surface in Fig.3B) would likely result in an association between the genotype and the phenotype. While true this assumption needs to be handed out carefully as it is not exhaustive. Indeed, some genetic paths may be highly structured and of relatively small amplitude. Examples of genetic path using real data from Dataset-1 will demonstrate this point.

### Examples of genetic path using the bone area of the ischium (BAI) as phenotype (Dataset-1)

The resulting average, ⟨θ_0_(i)⟩, and variance, σ(i), can be used to inform the null hypothesis of a particular SNP from ‘real’ datasets. However, since there are as many different sets of θ_0_(i)s as number of SNPs, each SNP will return its own ⟨θ_0_(i)⟩ (null hypothesis) upon scrambling. A comparison between SNPs using GIFT/genetic paths requires then to concentrate on the differences, Δθ(i) = θ(i) − ⟨θ_0_(i)⟩. In the remaining text one shall rewrite ⟨θ_0_(i)⟩ as θ_0_(i) to simplify notations.

Concentrating now on ‘real’ dataset, the genetic paths were obtained further to ranking BAI residual values (Dataset 1) using an incremental rank from small to large values. As an example, Fig.4 shows the two genetic paths θ(i) and θ_0_(i) for six SNPs, renamed SNP1-6 (see Table-2 for accurate genetic information) enabling us to appreciate the qualitative difference between the genetic paths. While the null hypothesis, i.e., θ_0_(i), resulting from the scrambling of phenotypic values many times always returns a straight line with a different slope for each SNP as seen above, the θ(i)s have different shape. To represent the set of θ(i)s in relation to the different microstates involved, each datapoint of the θ(i)s is colour coded as in Fig.1C.

**Figure 4:**
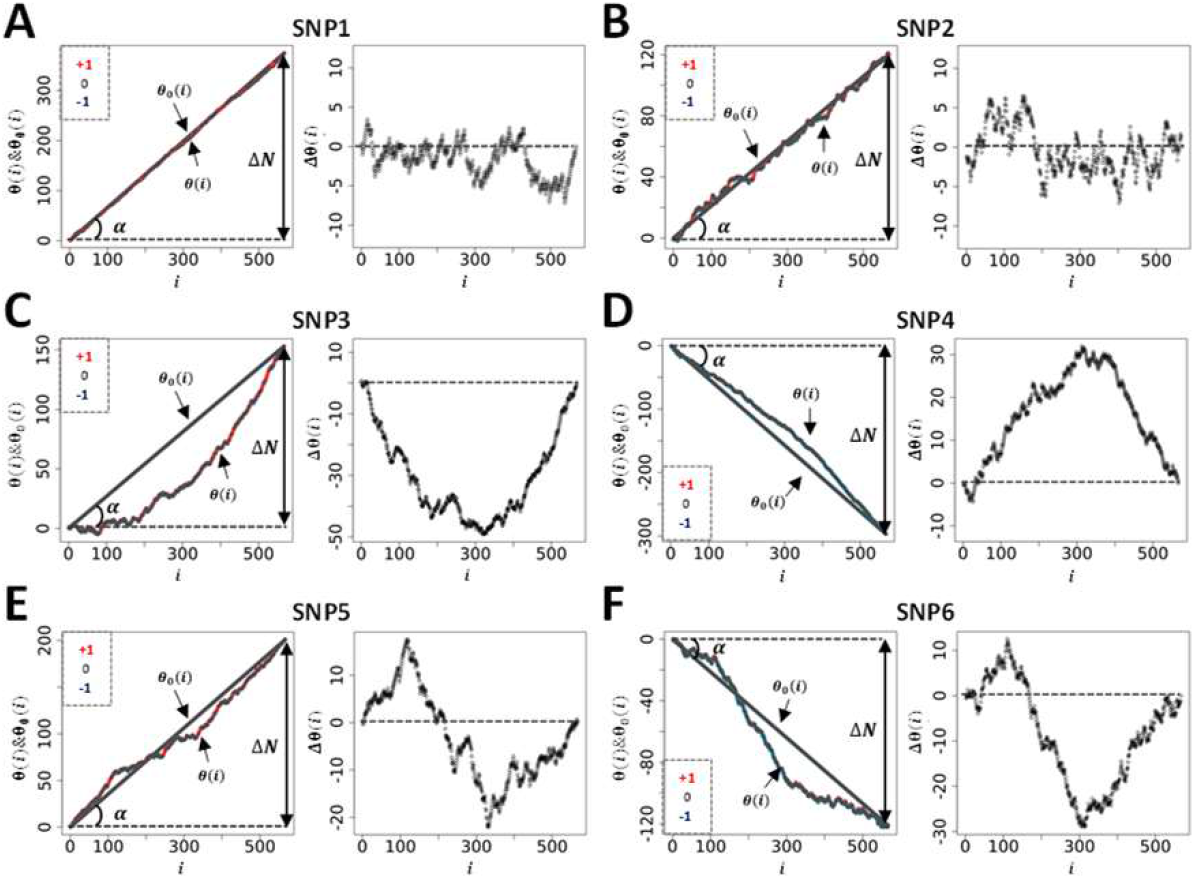
A sample of genetic paths selected from Dataset-1. The details of the different SNPs displayed are given in Table 2.

**Table 2:** Determination of gene/size effect (a) and dominance (d) for SNP1-6 from dataset-1. The level of significance for GIFT and GWAS is colour coded: red=not-significant, green=significant.

| CHR | NAME | POSITION | $-\text{Log}_{10}(p_{\text{GIFT}})$ | $-\text{Log}_{10}(p_{\text{GWAS}})$ | $N_+$ | $N_0$ | $N_-$ | a* | d** |
| --- | --- | --- | --- | --- | --- | --- | --- | --- | --- |
| 9 | OAR9_58767921 (SNP1) | 56039025 | 2.7895 | 0.2735 | 391 | 160 | 16 | N/A | N/A |
| 3 | s02120 (SNP2) | 213625709 | 2.8893 | 0.0018 | 198 | 291 | 78 | N/A | N/A |
| 6 | OAR6_40855809 (SNP3) | 36655091 | 28.5105 | 9.8639 | 229 | 262 | 76 | 96.85 | -13.01 |
| 6 | OAR6_38315830 (SNP4) | 34256151 | 20.7541 | 3.7366 | 24 | 222 | 321 | -70.02 | -0.05 |
| 23 | OAR23_35510473 (SNP5) | 33556377 | 19.7239 | 0.2301 | 254 | 260 | 53 | N/A | N/A |
| 25 | OAR25_30372586 (SNP6) | 29046746 | 18.5806 | 1.0692 | 90 | 266 | 211 | N/A | N/A |

(a\*): The gene/size effect is calculated considering the mid-distance between the average values of phenotypic residuals of microstates '-1' and '+1'. (d\*\*): The dominance is calculated considering the difference between the gene/size effect (a) and the position of the average value of phenotypic residuals for the microstate '0'.

Since θ_0_(i) is linked to the difference between the genetic microstate frequencies of homozygotes, ΔN = N_+_ − N_−_, in Fig.4 we represent by the angle α such difference. Since tan(α) = +N_+_/N − N_−_/N where N is the total number of positions (i = 1, 2, …, N) θ_0_(i) can be rewritten as, θ_0_(i) = tan(α)i. As any analysis must concentrate on the difference, Δθ(i) = θ(i) − θ_0_(i), such as to cancel the apparent variability in the null hypothesis across SNPs, we represent the plots of the different Δθ(i)s obtained in the right panel of Figs.4A-4F.

Figs.4A-4B display two distinct genetic paths that are globally similar. While they have different number of microstates of each type (see Table-2) the Δθ(i)s of SNP1 and SNP2 are characterized by their small amplitudes and the fact that they are erratic crossing several times the axis of position corresponding to the null hypothesis. In those cases, using the information contained in the phenotypic residuals, namely ranking the phenotypic residuals from small to large values, does not permit to fully differentiate θ(i) from θ_0_(i). On the other hand, the right panel in Figs.4C-4D for SNP3 and SNP4 demonstrates, in a more noticeable way, a paraboloid shape for the Δθ(i)s resulting from a segregation of microstates upon ordering the phenotypic residuals. The segregation of microstates +1 and -1 in opposite direction is reminiscent of Fisher theoretic works (Fig.1). As it turns out Figs.4C-4D show some similarities with Fig.1C based on a simulation inspired by Fisher’s seminal works. Importantly the ΔN-values of SNP1 and SNP4 while of opposite sign are similar in absolute value, are as those of SNP2 and SNP3, suggesting, in turn, the ΔN-values do not impact on the ability to differentiate θ(i) from θ_0_(i). Namely that a segregation of microstates can be inferred also with relatively large and opposed ΔN-values.

Envisaging the migration of microstates +1 and -1 in opposite direction as initially postulated by Fisher as the sole framework to associate genotype and phenotype is not always valid. This is demonstrated by SNP5 and SNP6 and the appearance of structured genetic paths displaying clear sigmoidal shapes for the Δθ(i)s as shown in Figs.4E-4F. Theoretically this phenomenon can be understood and explained by the presence of non-linear phenotypic fields, see (11) also reviewed in (12), in turn breaking the symmetry postulated by Fisher assuming the sole presence of linear phenotypic fields. This type of sigmoidal shapes is of interest since they inform on potential regulation mechanisms involving very probably ‘regulatory variants’ (21). Indeed, the right panels in Figs.4E-4F can be envisioned as representing the genetic organization of two distinct subpopulations of phenotypic residual values, one above the dashed line and the other one underneath it. Taken separately those two subpopulations draw curves like Figs.4C-4D or Fig.1C. In this context it is tempting to suggest that sigmoid genetic paths reveal a type of genotype-phenotype association that is inherently ‘scale-dependent’, namely function of the magnitude of phenotypic residuals. Because traditional GWAS concentrates on averages and variances, these sigmoid paths would be remarkably difficult to characterize with traditional methods. This is so because there is no clear antisymmetric segregation of microstates. As an example, using SNPs1-6 (from Fig.4) we have plotted, in Fig.5, the average values of phenotypic residuals for each microstate, and in Table-2 we provide the resulting gene/size effects and the dominances associated with those. Fig.5 and Table-2 demonstrate that sigmoid genetic paths (SNP5 and SNP6) are much less detectable with traditional methods while paraboloid genetic paths (SNP3 and SNP4) are. Note that the numerical determination of ‘−Log_10_(p_GIFT_)’ in Table-2, that is the significance for GIFT, is explained in the next part below.

**Figure 5:**
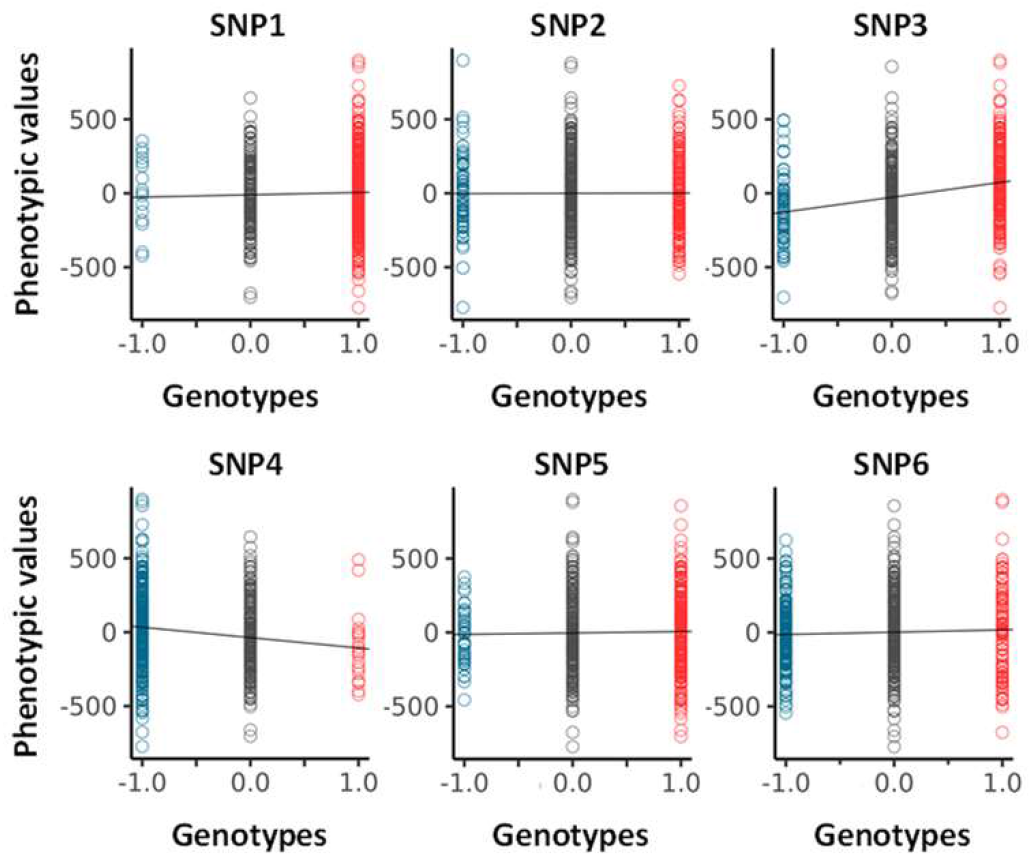
Analysis of averages (GWAS) for SNP1-6 (see Fig.4 and Table-2). Values for the size/gene effects (a) and dominances (d) are given in Table 2.

To conclude, based on Fisher’s theoretic works, the traditional GWAS method has been optimized to map SNPs that, using GIFT, would draw paraboloid genetic paths (see Fig.1C). The potential novelty using GIFT resides in its ability to provide new information and detect relatively regular/structured sigmoid genetic paths that would otherwise not be detected by traditional methods.

### P_GIFT_: p-value for GIFT

GIFT and GWAS extract information on genotype-phenotype associations in totally different ways. While GIFT concentrates on the significance of curves drawn using Δθ(i) = θ(i) − θ_0_(i), GWAS focuses solely on the significance of difference of averages. However, to compare GIFT to GWAS it is essential to determine a p-value for GIFT that is exhaustive enough such as to also capture the information that GWAS provides. To this end a p-value was derived that concentrates on the maximal amplitudes difference of genetic paths (see Figs.6A-6B).

The p-value for GIFT can be understood as follows. Since the number of possible paths is linked to the number of configuration possible resulting from lodging N_+_, N_0_ and N_−_ microstates into a list composed of N = N_+_ + N_0_ + N_−_ components, the number of possible paths is, 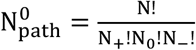. Let us now divide the genetic paths into regions, Δi_1_, Δi_2_ and Δi_3_ as shown in Figs.6A-6B. As the number of microstates of each sort can be determined in each region using an adequate algorithm, then the total number of possible genetic paths in this first, second and third regions are, respectively, 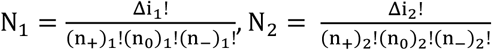 and, 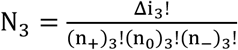, where (n_q_)_p_ is the number of microstate of type q in the p^th^ region, q ∈ {+,0, −} and p ∈ {1,2,3}. Consequently, the probability of a genetic path in this context is, 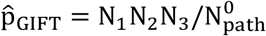. Using the null hypothesis simulations shown in Fig.3 based on the theoretic SNPs given in Table-1, 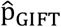 may be determined for each genetic path simulated. Its statistic plotted in Fig.6C for each SNP demonstrates very little variations across SNPs or when the sample size changes by a factor two. Based on this observation confidence intervals were determined for all SNPs by averaging the 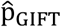 values obtained. The upper and lower red dashed lines represent the 99% and 95% confidence intervals. To consider the false discovery rate (FDR) and adjust p-values to remove type-I errors, 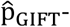 values in Fig.6C were corrected using the Benjamini-Hochberg procedure leading to a new set of adjusted, i.e., reduced, p-values, noted p_GIFT_ (see Fig.6D), that may be used to determine the true significance of DNA variants (SNPs). Returning to Table-2 the numerical value of p_GIFT_ was determined for the genetic paths shown in Fig.4 demonstrating that GIFT can extract information when sigmoid genetic paths are involved while traditional GWAS is unable to do so.

**Figure 6:**
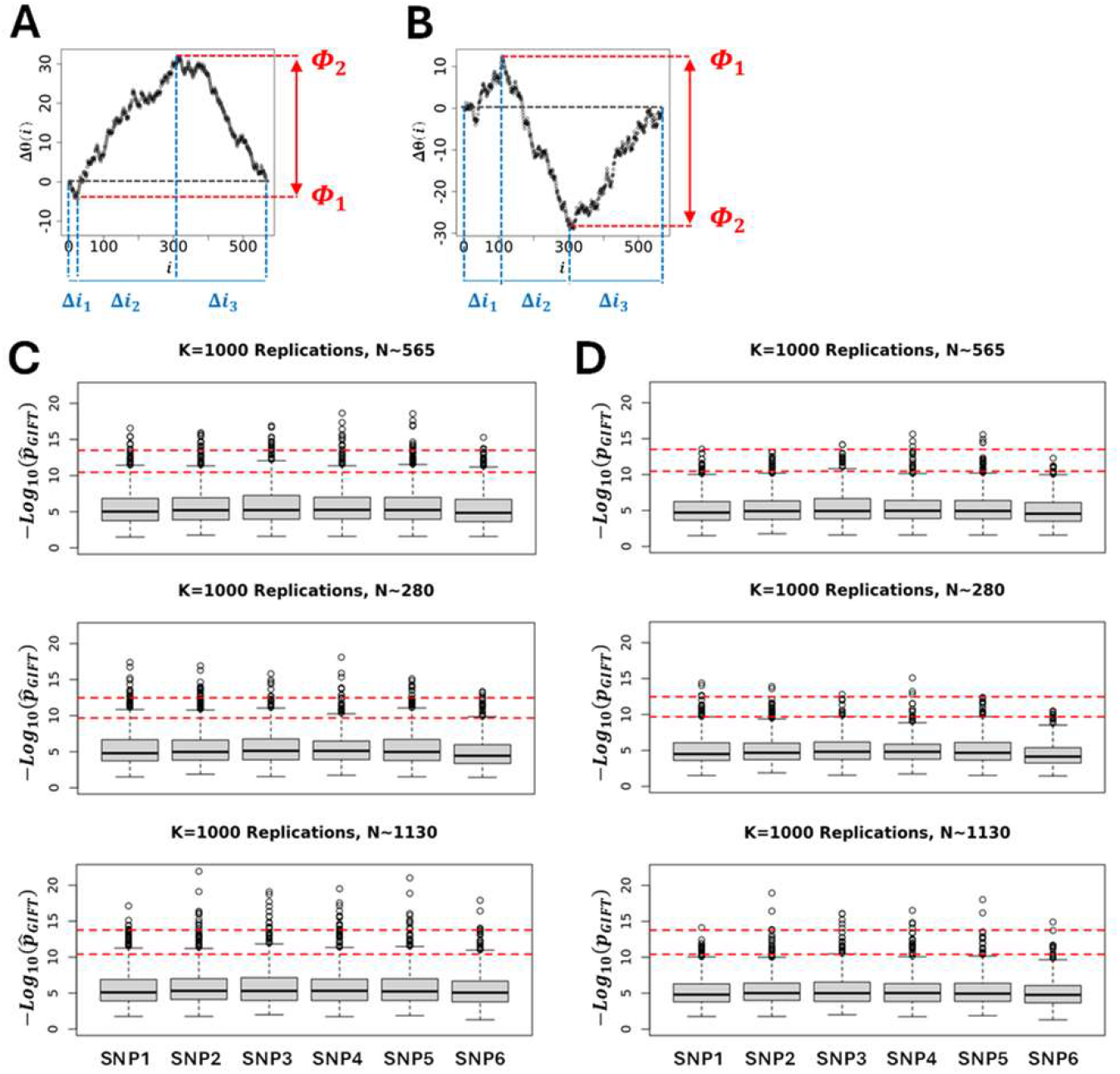
To provide a p-value extracting genotype-phenotype associations in an exhaustive manner for both GWAS and GIFT a method concentrating on the largest and smallest extreme values of the genetic path was focused upon. This method can be applied to paraboloid (GWAS or GIFT-like) **(A)** and sigmoid (GIFT-like) **(B)** genetic paths. The overall idea consists in determining how many paths *N*_1_, *N*_2_ and *N*_3_ can be generated from the respective interval of positions Δ*i*_1_, Δ*i*_2_ and Δ*i*_3_ given that the constraints for the extrema are Φ_1_ and Φ_2_. Then a p-value 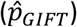 can be determined as seen in the text. **(C)** Using simulations (K=1000 replicates) a statistic of 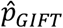 for the null hypothesis can be generated using theoretic SNPs (Table-1). Simulations demonstrate that 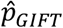 is relatively independent of the microstate’s frequencies upon which a 99% (upper dashed line) and 95% (lower dashed line) interval confidences can be generated. **(D)** 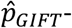-values were adjusted to consider FDR using Benjamini-Hochberg procedure leading to a new set of *p*_GIFT_-values. The code for the simulations is given in Supplemental S4.

Armed with p_GIFT_ an analysis of datasets can now be performed.

### Comparison between GWAS and GIFT considering the bone area of the ischium (BAI) as phenotype (Dataset-1)

The first dataset (Dataset-1) analysed 567 pedigree-recorded Scottish Blackface lambs concentrating on the bone areas of the ischium measured in mm^2^ from cross-sectional CT scans (13). After adjusting phenotypic values, the work demonstrated a clear involvement of chromosome 6 as shown in Fig.7A. The genome-wide significant thresholds applied for GWAS in Fig.7A correspond to Bonferroni corrections at 1% (upper red dashed line) and 5% (lower dashed red line) determined by using independent SNPs only. Formally a 1% (resp. 5%) Bonferroni correction is given by, −Log_10_(0.01/N_ind−SNPs_) (resp. −Log_10_(0.05/N_ind−SNPs_) where N_ind−SNPs_ = 10433 is the number of independent SNPs. Using its own thresholds (Fig.6D) GIFT was applied using the same set of phenotypic residuals. Figs.7A-7B demonstrate the results obtained by GWAS and GIFT using Manhattan plots.

**Figure 7:**
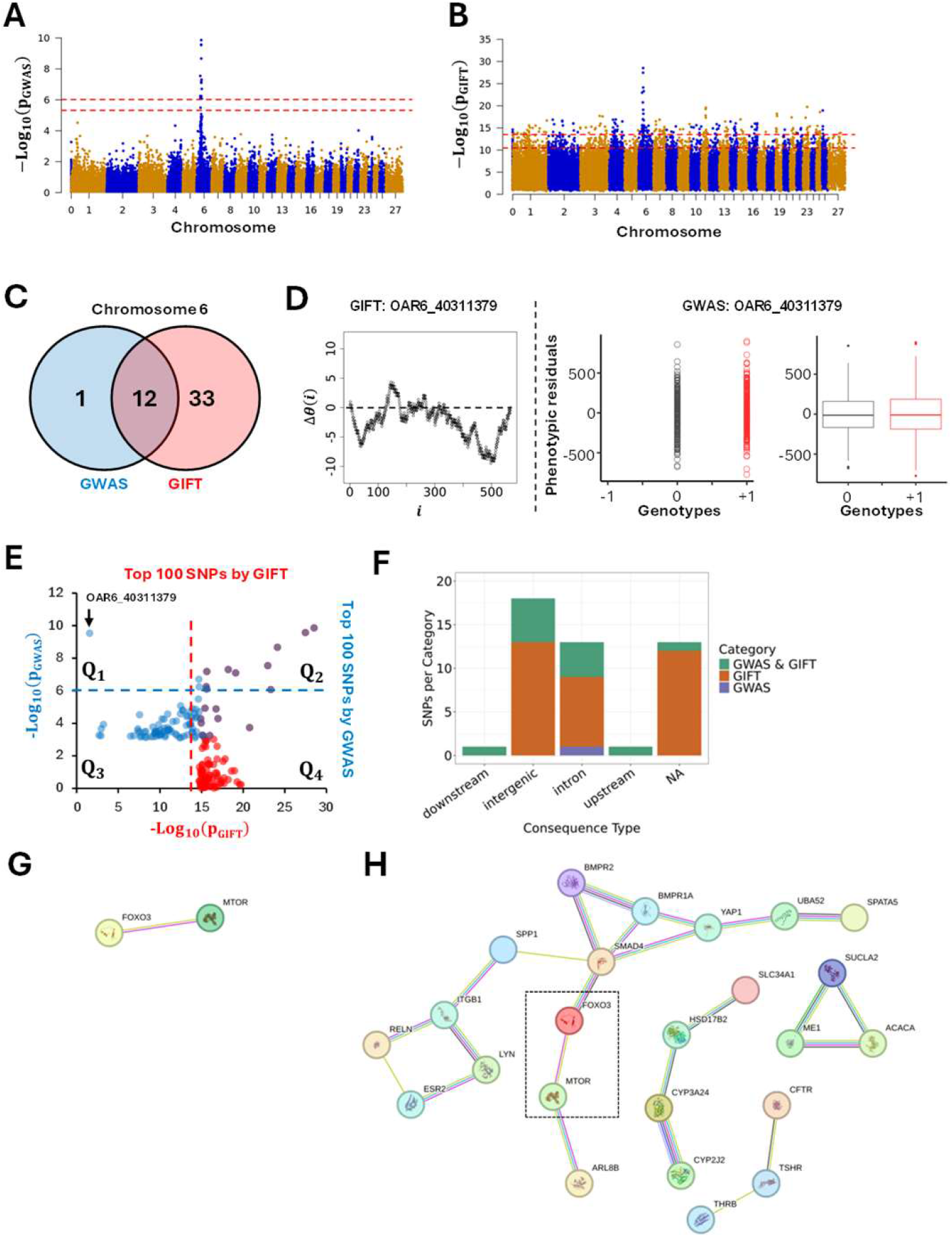
Manhattan plots based on p-values obtained by GWAS **(A)** and GIFT **(B)** demonstrating significant differences between the methods concerning potential genotype-phenotype associations. Note that the presence of a chromosome ‘0’ results from the fact that some SNPs identified by (Matika et al., 2016) were not allocated to specific chromosomes/genomic positions due to lack of information at the time. A fathom chromosome (chromosome zero) was created to allocate those SNPs. **(C)** Venn-diagram representing the most significant SNPs by GWAS and GIFT. One SNP (OAR6_40311379) demonstrated a large p-value for GWAS and a small p-value for GIFT. A representation of its genetic path **(D-left)** did not underscore any ‘parabolic’ or ‘sigmoidal’ associations. As it turned out this SNP was a false-positive by GWAS since the difference between the phenotypic means was not significative **(D-right). (E)** The 100 most significant SNPs by GWAS and GIFT were extracted, and their p-values plotted against each other. The dashed lines represent the threshold applied for GWAS (blue dashed line) and GIFT (red dashed line). The SNP OAR6_40311379 pointed by the black arrow is the single one standing out in *Q*_1_ confirming its false-positive status. **(F)** Biotypes of the most significant SNPs by GIFT and GWAS. **(G)** String analysis performed to determine gene networks using significant SNPs by GWAS. **(H)** String analysis performed to determine gene networks using significant SNPs by GIFT, note that the dashed square underlines mTOR and FOXO3 determined by GWAS. The code for obtaining Figs.7B, 7C, 7D, 7F is given in Supplemental S7.

The significance threshold by GIFT was defined by a null hypothesis using theoretic SNPs. To demonstrate that the theoretic results obtained from Fig.6D are transferrable to ‘real’ SNPs (Fig.7B), namely that the significant SNPs obtained in Fig.7B have null hypotheses with similar properties like those shown in Fig.6D, each significant SNP (Fig.7B) had its genetic path randomly permutated a thousand times to determine the distribution of −Log_10_(p_GIFT_)-values corresponding to their null hypothesis. Results show that the null hypotheses are remarkably similar across SNPs and that the threshold determined using theoretic SNPs (Fig.6D) holds when ‘real’ SNPs are used (Supplemental S5).

Overall, Fig.7A and Fig.7B demonstrate that there is an agreement between GWAS and GIFT that chromosome 6 is involved. However, differences exist that are shown through the involvement of several chromosomes when GIFT is used. Considering the thresholds involved, for GWAS the phenotype studied may be considered as a sort of ‘single gene trait’ while for GIFT, the phenotype looks very much like a ‘complex trait’ involving more chromosomes than chromosome 6. Detailed information of all significant SNPs by GWAS or GIFT is given in Supplemental S6.

Concentrating on Chromosome 6 to address the overlap of information provided by GIFT and GWAS, a Venn-diagram including highly significant SNPs only, namely SNPs beyond the upper red dashed-line in Figs.7A-7B, was plotted. The Venn-diagram (Fig.7C) reveals that most SNPs deemed significant by GWAS were also deemed significant by GIFT. Curiously, only one SNP seemed highly significant by GWAS but irrelevant for GIFT. As p_GIFT_ was designed to collect exhaustive information from GWAS, the SNP was identified (OAR6_40311379) and its genetic path, i.e., its Δθ(i), plotted (Fig.7D-left) together with its GWAS-representations (Fig.7D-right). The genetic path, being erratic of relatively small amplitude and crossing several times the axis of positions, did not display any obvious ‘parabolic or sigmoidal’ associations at first sight, in turn justifying its small p_GIFT_-value. The GWAS-representation of OAR6_40311379 however, demonstrated the absence of microstate ‘-1’ as well as a near overlap of microstates ‘0’ and ‘+1’ further demonstrated by the similarities between their boxplots, suggesting the occurrence of a false-positive. To confirm this a comparison of phenotypic means for the microstates ‘0’ and ‘+1’ was performed returning a t-test value of 1.1485 (p-value of 0.2512), confirming the presence of a false-positive.

In order to assess the overlap of information between GWAS and GIFT we plotted in Fig.7E the first 100 more significant SNPs detected by GIFT and GWAS. Results confirm an overlap of SNPs associated with the phenotypic residuals for large values of p_GIFT_ and p_GWAS_ (see purple dots in Q_2_ in Fig.7E). Interestingly, two SNPs considered as significant by GWAS (two blue dots in Q_2_) were not by GIFT. That is because the p_GIFT_-values for these dots were less than other SNPs detected by GIFT. As already stated above many SNPs from other chromosomes were considered significant by GIFT that were not by GWAS (see red dots in Q_4_). Finally, the quadrant Q_1_ in Fig.7E confirms that OAR6_40311379, i.e., the false positive detected by GWAS, is a standalone SNP among the 100 SNPs for which p_GWAS_ > p_GIFT_. Finally, the biotype of significant SNPs on Chromosome 6 for GIFT and GWAS are also presented in Fig.7F.

The primary conclusion provided by Figs.7A-F is that, when compared to GWAS, GIFT returns substantially more genetic information.

However, a central question concerns the genetic pertinence of the significant SNPs obtained by GIFT. As GIFT has been designed with the aim to increase the investigative power of biological datasets, we may assume that the significant SNPs obtained by GIFT once translated into gene names should underline some level of non-random gene-gene interactions. The latter point is particularly relevant since GIFT is expected to detect regulatory variants (c.f. sigmoidal genetic paths). To assess this point we performed an enrichment analysis based on gene names using the String database, which helps determine known and predicted protein-protein interactions. In order to apply String the significant SNPs obtained using GWAS and GIFT were mapped to the reference sheep genome assembly from ensembl (Oar_v3.1) to obtain the gene names. Using those gene names String analyses were performed for GWAS and GIFT using a minimum required interaction score of 0.4. Fig.7G and Fig.7H show the networks obtained. With enrichment p-values for GWAS and GIFT of 0.176 and 0.00008, respectively, these results confirm that the set of genes determined by GIFT have more interactions among themselves than what would be expected for a random set of genes of the same size and degree distribution drawn from the genome. Namely that GIFT increases the investigative power of biological datasets.

At present, we do not know how the whole information provided by GIFT may inform on the putative biology of the phenotype studied (BAI). As it turns out, a full validation of the information provided by GIFT on Dataset-1 would require an in-depth mutational/deletion/insertion/gene-editing analyses in live animals, extending beyond the scope of this present article.

To demonstrate the relevance of the information provided by GIFT we decided to challenge GIFT using a different dataset (Dataset-2) concentrating on a complex trait related to 1C-metabolism.

### Comparison between GWAS and GIFT considering 1C-metabolites as phenotype (Dataset-2)

Dataset-2 concerns biochemical data which seeks to identify risk allele variants in genes whose products direct a specific series of metabolic pathways, known as one carbon (1C) metabolism (Fig.2). The significance of 1C metabolism is that it is a complex trait involving a series of interlinking metabolic pathways that provide 1C units (methyl groups) for the synthesis and methylation of biological molecules. After 1% and 5% Bonferroni corrections for GWAS and the Benjamini-Hochberg procedure applied to GIFT, the Manhattan plots were obtained (Fig.8A). Note that the number of independent SNPs in this case is 624 (out of 3923 SNPs from the gene array). Fig.8A demonstrates clearly that the informational power of GWAS is less than that of GIFT. Finally, in Fig.8B we provide the biotypes of the most significant SNPs shown by the upper red dashed lines obtained using GIFT. Detailed genetic information of the most significant SNPs obtained using GIFT is provided in Supplemental S8.

**Figure 8:**
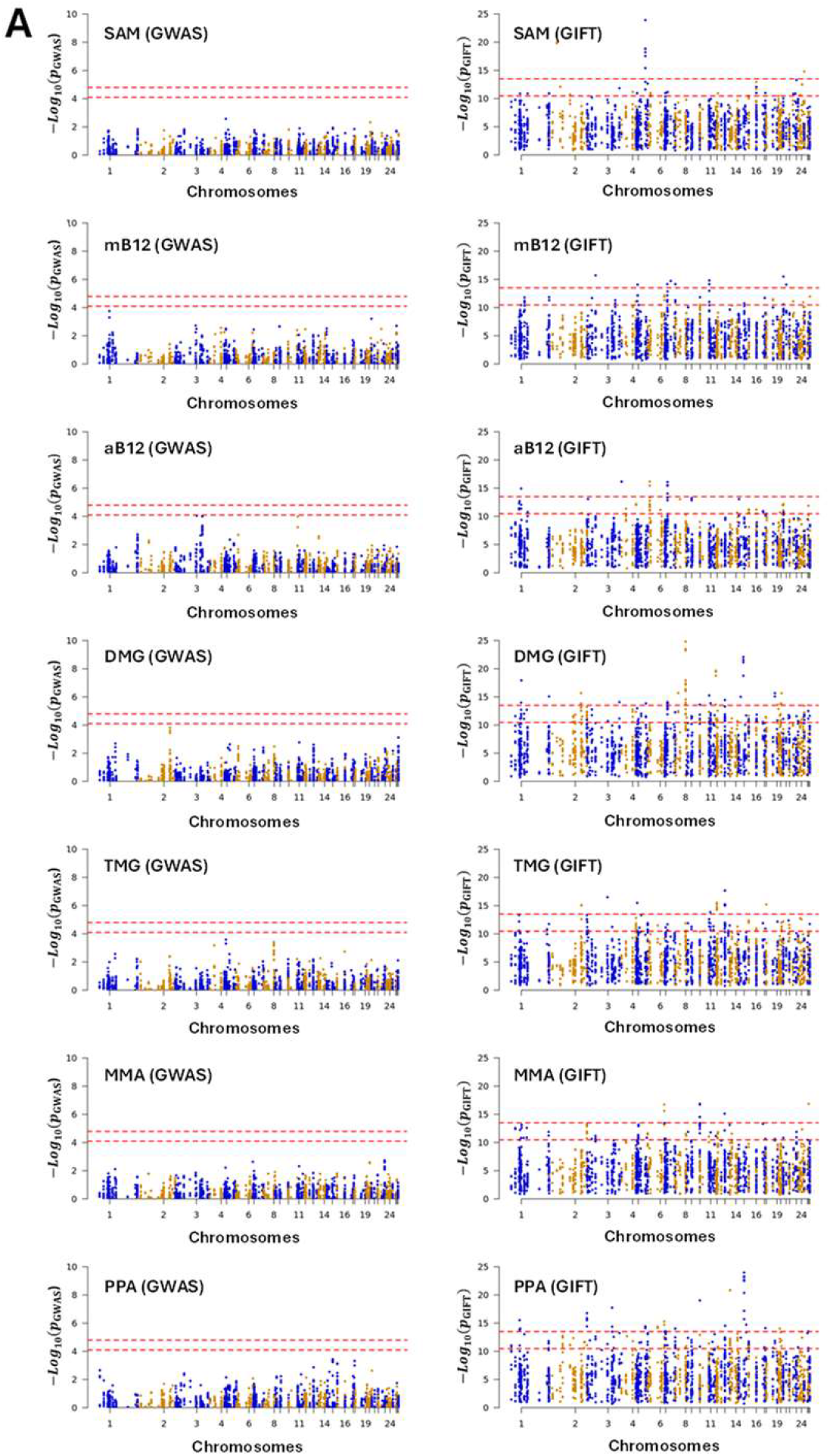
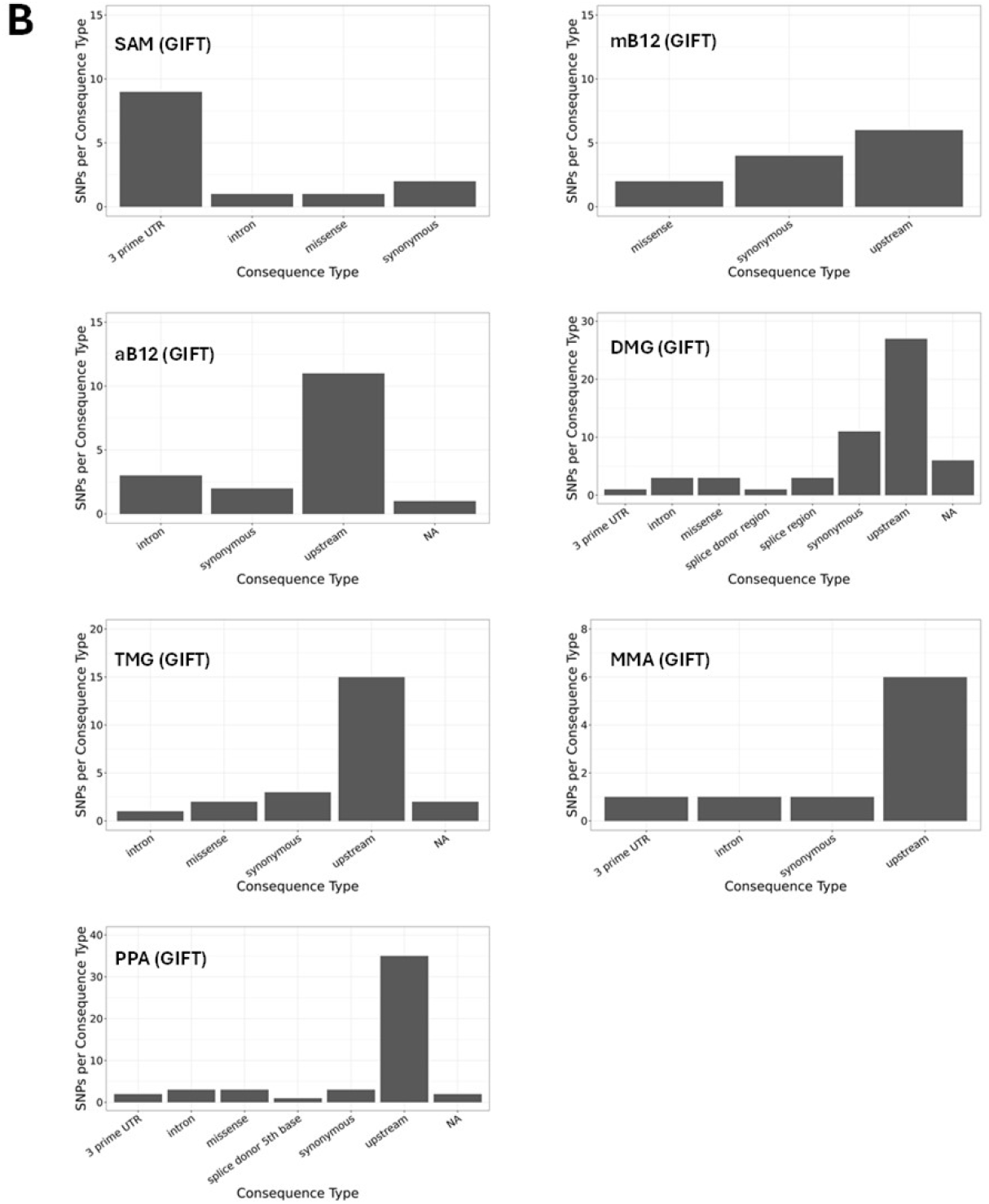
**(A)** Comparison of the information extracted by GWAS and GIFT using Manhattan plots for the metabolites presented in red in Fig.2. We recall the acronyms, S-adenosyl methionine (SAM), methylcobalamin (mB12), adenosylcobalamin (aB12), trimethylglycine (TMG), dimethylglycine (DMG), propionate (PPA) and methylmalonic acid (MMA). It should be noted that due to inherent difficulty linked to the measure of metabolite the sample sizes were not similar across metabolites, that is the values for N differ between the Manhattan plots (SAM: N=344; mB12: N=183; aB12: N=338; DMG: N=338; TMG: N=340; MMA: N=348; PPA: N=345). **(B)** Biotypes corresponding to the most significant SNPs for each metabolite determined by GIFT (a detailed list of information concerning those SNPs in given in supplemental S8). The code for the Manhattan plots and the determination of biotypes is given in Supplemental S9.

Since the gene array was synthesized using SNPs from known genes involved in 1C metabolism, the relevance of String analyses (i.e., enrichment p-values) would be minimal and of little interest.

Besides validating that GIFT may extract more information from genotype-phenotype datasets, it is worth underlying the biological importance and novelty of results obtained. One carbon metabolism in sheep is comparable to that in humans. The significance of 1C metabolism is that it is a complex trait involving a series of interlinking metabolic pathways that provide 1C units (methyl groups) for the synthesis and methylation of chromatin among other molecules (15). S-adenosylmethionine (SAM) is a potent methyl donor within these cycles and serves as the principal substrate for methylation of DNA, associated proteins, and RNA. It was previously demonstrated in sheep, cattle, rodent and human studies that disrupting these cycles during early pregnancy, by either dietary means (i.e., reducing dietary vitamin B12, folate, choline and/or methionine), or through exposure to environmental chemicals such as cigarette smoking, can lead to epigenetic dysregulation and impaired foetal development with long-term consequences for offspring cardiometabolic health (22–25). It was also advocated that interindividual and ethnic variability in epigenetic gene regulation arises because of single-nucleotide polymorphisms (SNPs) within 1C genes, associated epigenetic regulators, and differentially methylated target DNA sequences (15). However, information concerning the nature and extent of interactions between parental genotype, diet and EC exposure was, until now, limited to just a few 1C genes in humans (15). Consequently, data obtained by the current study provide new evidence concerning significant genetic variants in 1C-metabolism and directly associated metabolic genes and epigenetic regulators that rely on SAM as the methyl donor, potentially applicable to the human species.

## DISCUSSION

While statistical association methods should not favor any biases when analyzing datasets, the way they are built mathematically is often indicative of a particular way of thinking. For example, with GWAS the phenotype is decomposed onto more fundamental sub-distributions characterized by the distribution of microstates (see Fig.1A). This approach underlines a sort of bottom-up approach that, within a reductionist framework, defines genes as biological agents controlling the phenotype aligned with the ‘Neo-Darwinian synthesis’. However, nothing prevents considering the opposite as far as statistical association methods are involved, and GIFT uses this degree of freedom. By using the full range of phenotypic information, GIFT transforms a random or disordered string of microstates (the straight line in the asymptotic limit seen in Fig.1C or Fig.3A) into an ‘ordered’ configuration of microstates (see Fig1C or Figs.4C-4F), in turn providing the signature of a genotype-phenotype association. Accordingly, since the phenotypic information controls the configuration of microstates it is a top-down approach, which turns out to be remarkably sensitive. GIFT has been estimated to be ∼1000 more sensitive than GWAS (11).

There are three main reasons as to why GIFT is more sensitive. The first is that GIFT determines the significance of curves composed of an entire population of datapoints. As curves provide a greater level of significance than considering differences between microstate/phenotypic averages/variances as advocated by GWAS, hence GIFT is statistically more powerful. The second reason is that the null hypothesis for GIFT, namely θ_0_(i), is contained in the definition of Δθ(i) and is therefore specific to the genome position, or SNP, studied. With GIFT there are as many null hypotheses as SNPs. This contrasts with GWAS defining a null-hypothesis valid for all SNPs at the population level when the average of microstate distributions overlap. Consequently, the discriminative power of GIFT is amplified. The third reason is that GIFT is simpler than GWAS. Indeed, based on R.A. Fisher’s seminal work, GWAS is based on a complex theory that seeks to determine genotype-phenotype associations on one hand (aim 1), and the heritability of phenotypes/traits studied on the other (aim 2). To achieve those two aims, the GWAS approach relies on frequentist probability to determine the validity of statistical inferences giving the notions of average and variance fundamental meanings related to aim 1 and 2, respectively. However, because average and variance are antinomic it is nearly impossible to have a clear picture of associations (size effects) since the noise (variance/heredity) blurs the average(s). On the other hand, by concentrating on genetic paths (curves) GIFT determines a global association. This does not mean that GIFT rules out the notions of size effect, dominance, and heritability, on the contrary, it encapsulates them under the generic notion of phenotypic field, i.e., size effect, dominance and heritability can be rederived from the phenotypic field. The term ‘field’ in the acronym GIFT is used to explain the disorder-order transition in the string of microstates using an analogy related to physics field theory, see (11, 12) for more details.

Finally, it is important to reframe GIFT within current debates in the field of biology. With GIFT it is the (information on the) phenotype that selects which SNP is required for its subsistence and it is interesting to note that, at the conceptual level and as a top-down approach, GIFT has some familiarity with the notion of phenotypic plasticity. Phenotypic plasticity refers to the ability of phenotypes to respond to a change in the environment favoring a divergence from the ancestor phenotype. As the phenotype relies on traits (modules), the responsiveness to any new input(s) must involve a re-organization of the phenotype architecture by allowing phenotypic sub-components (modular traits) to adapt the changes (26). Namely that genetic accommodation linked to a standing pool of genetic variations characterizing any trait is central to phenotypic plasticity that, through persistence, may genetically assimilate the new architecture (selection) (26, 27). In this context the top-down method GIFT, which is essentially a phenotype-genotype (and not genotype-phenotype) association method, can pull out any standing genes awaiting to be used by phenotypes.

To conclude, we provide evidence that GIFT enhances the investigative power of biological datasets. Additionally, we provide evidence also for the need to rethink the conceptual bases of genotype-phenotype association methods, such as use more information from the whole biodiversity of data.

## DATA AVAILABILITY

Data including supplementary materials are available using the link:

## SUPPLEMENTAL MATERIAL

S1 provides the raw data for Dataset-1; S2 provides the raw data for Dataset-2. S3 provides the statistical summary for the phenotypic adjustment prior to running GWAS on Dataset-2; S4 provides the code to obtain Fig.3 and Fig.6C. S5 represents the permutation analysis of significant SNPs obtained by GIFT from dataset-1. S6 represents the list of significant SNPs obtained by GWAS and GIFT when applied on Dataset-1; S7 provides the code to obtain Fig.4, Fig5 and Fig.7. S8 provides the list of significant SNPs by GIFT for Dataset-2. S9 provides the code to obtain Fig.8.

## ACKNOWLEDGMENTS

The authors would like to thank Dr Barbara Bravi, Dr Wing-Yee Kwong and Dr Dongfang Li for useful discussions and/or technical assistance.

Present address of B.B.: Imperial College, London, UK.

Present address of W-Y.K. & D.L.: University of Nottingham, Sutton Bonington, UK.

## AUTHOR CONTRIBUTIONS

CR conceptualized GIFT; CR & JW formalized GIFT; PK coded GIFT simulations; Dataset-2 was designed and obtained by KDS, OM, AHB, CEC, JX, DAB, RDE; Dataset-1 and Dataset-2 were analysed by PK, CR, JW, KDS, OM, AP; paper was written by CR and KDS, and proofread by CR and KDS.

