## Supplementary material for "Investigative power of Genomic Informational Field Theory (GIFT) relative to GWAS for genotype-phenotype mapping": S4

Simulation of null hypothesis


### Simulation of null hypothesis

###### Supplemental material by Kyratzi et al.

First, we build SNPs with the same sample size of \(N=565\), by fixing the number of ‘+1’
states and by transferring 100 microstates from ‘-1’ to ‘0’. We then
shuffle this list multiple times \((K=1000)\) and run GIFT on these lists to
generate theta paths. Subsequently, we calculate the standard deviation
of \(\Delta \Theta/\sqrt N\) values and
generate plots against position \(j/N\). Also, we calculate the \(pGIFT\) and generate the boxplots for each
SNP per replication \(K\).

We perform similar analyses varying a proportion of \(N\_+, N\_-, N\_0\) to get half (\(N\sim 280\)) and double (\(N\sim 1130\)) the sample size.

### 1. Set the functions needed to run GIFT

```
# Function to compute the cumulative distributions of the ordered and random configurations
# Input variables:
#   Geno: sorted microstates dataset (row: individuals, col: SNPs)
#   Nmpz: dataframe of number of microstates for each SNP (rows)
# Output variables:
#   List of Wp, Wm, Wz, W0p, W0m, W0z: Cumulative distributions of +1,-1 respectively for ordered and random configurations

Fun_CumDistr <- function(Geno, Nmpz){
  library(tidyr) # tibble, pivot_wider
  library(dplyr)
  library(zoo) # na.locf
  
  CumDistrW <- as.data.frame(apply(Geno, 2, function(x){
    # Convert vector of microstates to data frame
    df <- tibble(position_j = seq_along(x), value = x)
    uniq.val <- as.character(unlist(unique(df[!is.na(df[,2]),2])))
    
    # Create cumulative sum for each unique value
    result <- df %>%
      group_by(value) %>%
      mutate(cumulative_sum = cumsum(value == value)) %>%
      ungroup()
    
    # Pivot the result to have one column per unique value
    result <- pivot_wider(result, names_from = value, values_from = cumulative_sum, values_fill = 0)
    
    # Replace zeros with NA values from the index onward
    result <- apply(result[,uniq.val], 2, function(x){
      first_non_zero_index <- which(x != 0)[1]
      x[first_non_zero_index:length(x)][x[first_non_zero_index:length(x)] == 0] <- NA
      return(x)
    })
    
    # Use na.locf() to fill NA values with the last non-NA value
    result <- as.data.frame(na.locf(result))
    Wm <- unlist(result[,which(uniq.val=="-1")])
    Wp <- unlist(result[,which(uniq.val=="1")])
    Wz <- unlist(result[,which(uniq.val=="0")])
    
    # If have less than three states, replace the null vectors with zeros
    if (is.null(Wm)) {Wm <- rep(0, times = nrow(df))}
    if (is.null(Wp)) {Wp <- rep(0, times = nrow(df))}
    if (is.null(Wz)) {Wz <- rep(0, times = nrow(df))}
    
    return(list(Wp = Wp, Wm = Wm, Wz = Wz))
  }))
  
  Wp <- CumDistrW[,grep("Wp$", names(CumDistrW), value = TRUE)]
  Wm <- CumDistrW[,grep("Wm$", names(CumDistrW), value = TRUE)]
  Wz <- CumDistrW[,grep("Wz$", names(CumDistrW), value = TRUE)]
  
  N <- nrow(Geno)
  W0p <- c(1:N) * Nmpz[1,2] / N
  W0m <- c(1:N) * Nmpz[1,1] / N
  W0z <- c(1:N) * Nmpz[1,3] / N
  
  return(list(Wp = Wp, Wm = Wm, Wz = Wz, W0p = W0p, W0m = W0m, W0z = W0z))
}

# Function to calculate Theta-paths
# Input variables:
#   CumDistr: List consisting of the Cumulative Distributions Wp, Wm, Wz, W0p, W0m, W0z
# Output variables:
#   List of Thp = Wp - W0p,
#           Thm = Wm - W0m,
#           Thj = Wp - Wm (the phenotype-responding genetic path),
#           Th0j = W0p - W0m (the default genetic path),
#           DThj = Thj - Th0j

Fun_ThPaths <- function(CumDistr){
  
  N <- nrow(CumDistr$Wp)
  num_SNPs <- ncol(CumDistr$Wp)
  
  Thp <- data.frame(matrix(NA, nrow = N, ncol = num_SNPs))
  Thm <- data.frame(matrix(NA, nrow = N, ncol = num_SNPs))
  Thj <- data.frame(matrix(NA, nrow = N, ncol = num_SNPs))
  Th0j <- data.frame(matrix(NA, nrow = N, ncol = 1))
  DThj <- data.frame(matrix(NA, nrow = N, ncol = num_SNPs))
  
  Thp <- CumDistr$Wp - CumDistr$W0p
  Thm <- CumDistr$Wm -CumDistr$W0m
  Thj <- CumDistr$Wp - CumDistr$Wm
  Th0j <- CumDistr$W0p - CumDistr$W0m 
  DThj <- Thj - Th0j
  
  return(list(Thp = Thp, Thm = Thm, Thj = Thj, Th0j = Th0j, DThj = DThj))
}
```

#### Define pGIFT for three states

\[pGIFT=
\Bigg(\frac{1}{2}-\frac{1}{\pi}tan^{-1} \Big(\frac{\sqrt{-\phi
\tilde\phi}}{(\phi-\tilde\phi) \sqrt{2}} \Big) \Bigg) \exp\Big\{
-\frac{8N(\phi-\tilde\phi)^2}{N(N-N\_0)-(N\_+-N\_-)^2} \Big\}, \quad \phi =
\max\_{j}\Delta\Theta(j) , \quad \tilde\phi = \min\_{j}\Delta\Theta(j)
\]

```
# Function to calculate the pGIFT
# Input variables:
#   Geno: sorted microstates dataset (row: individuals, col: SNPs)
#   Nmpz: dataframe of number of microstates for each SNP
#   ThPaths: List of Theta-paths (output of Fun_ThPaths)
# Output variables:
#   pGIFT: vector consists of pGIFT values
Fun_pGIFT <- function(Geno, Nmpz, ThPaths){
  pGIFT <- c()
  
  num_SNPs <- ncol(Geno)
  DThj <- as.data.frame(ThPaths$DThj)
  
  phi <- apply(DThj, 2, function(x){max(x, na.rm = TRUE)})
  phi_tilde <- apply(DThj, 2, function(x){min(x, na.rm = TRUE)})
  
  for (i in 1:num_SNPs) {
    
    no.NA <- !is.na(Geno[,i])
    N <- sum(no.NA)
    
    e <- exp( -(8*N*(phi[i]-phi_tilde[i])^2) / (N*(N-Nmpz[i,3])-(Nmpz[i,2]-Nmpz[i,1])^2) )
    pGIFT[i] <- ( (1/2)-(1/pi)*atan(sqrt(-phi[i]*phi_tilde[i])/((phi[i]-phi_tilde[i])*sqrt(2))) ) *e
    
  }
  
  return(pGIFT)
  
}
```

### 2. Run GIFT for simulated data when microstates are shuffled \(K\) times.

#### \(K=1000, N \sim 565\)

```
# Define the dataset 
Nmpz <- rbind(data.frame(Nm = 515, Np = 25, Nz = 25), data.frame(Nm = 415, Np = 25, Nz = 125),
              data.frame(Nm = 315, Np = 25, Nz = 225), data.frame(Nm = 215, Np = 25, Nz = 325),
              data.frame(Nm = 115, Np = 25, Nz = 425), data.frame(Nm = 15, Np = 25, Nz = 525))

num_SNPs <- nrow(Nmpz)
rownames(Nmpz) <- paste("SNP", c(1:num_SNPs))
Nmpz
```

```
# Generate microstates dataset
K <- 1000
GenoRep1000 <- vector("list", num_SNPs)

for (i in 1:num_SNPs){
  set.seed(i)
  GenoRep1000[[i]] <- data.frame(replicate(K, sample(rep(c(-1,+1,0), times = c(Nmpz$Nm[i],Nmpz$Np[i],Nmpz$Nz[i]))), simplify = FALSE))
  colnames(GenoRep1000[[i]]) <- paste0("Rep",c(1:K))
}

# Call Fun_CumDistr
CumDistrRep1000 <- vector("list", num_SNPs)
for (i in 1:num_SNPs) {
  CumDistrRep1000[[i]] <- Fun_CumDistr(GenoRep1000[[i]], Nmpz[i,])
}

# Call Fun_ThPaths
ThPathsRep1000 <- vector("list", num_SNPs)
for (i in 1:num_SNPs){
  ThPathsRep1000[[i]] <- Fun_ThPaths(CumDistrRep1000[[i]])
}

# Call Fun_pGIFT
pGIFTRep1000 <- vector("list", num_SNPs)
for (i in 1:num_SNPs){
  pGIFTRep1000[[i]] <- Fun_pGIFT(GenoRep1000[[i]], Nmpz[i,], ThPathsRep1000[[i]])
}
```

```
# Calculate sd of DeltaTheta(j) for each position per replication
sdDThjRep1000 <- lapply(ThPathsRep1000, function(ThPaths){apply(ThPaths$DThj, 1, sd)})
```

#### \(K=1000, N \sim 280\)

```
# Define the dataset 
NmpzHalf <- floor(Nmpz/2)
num_SNPs <- nrow(NmpzHalf)
rownames(NmpzHalf) <- rownames(Nmpz)
NmpzHalf
```

```
# Generate microstates dataset
K <- 1000
GenoRep1000Half <- vector("list", num_SNPs)

for (i in 1:num_SNPs){
  set.seed(i)
  GenoRep1000Half[[i]] <- data.frame(replicate(K, sample(rep(c(-1,+1,0), times = c(NmpzHalf$Nm[i],NmpzHalf$Np[i],NmpzHalf$Nz[i]))), simplify = FALSE))
  colnames(GenoRep1000Half[[i]]) <- paste0("Rep",c(1:K))
}

# Call Fun_CumDistr
CumDistrRep1000Half <- vector("list", num_SNPs)
for (i in 1:num_SNPs) {
  CumDistrRep1000Half[[i]] <- Fun_CumDistr(GenoRep1000Half[[i]], NmpzHalf[i,])
}

# Call Fun_ThPaths
ThPathsRep1000Half <- vector("list", num_SNPs)
for (i in 1:num_SNPs){
  ThPathsRep1000Half[[i]] <- Fun_ThPaths(CumDistrRep1000Half[[i]])
}

# Call Fun_pGIFT
pGIFTRep1000Half <- vector("list", num_SNPs)
for (i in 1:num_SNPs){
  pGIFTRep1000Half[[i]] <- Fun_pGIFT(GenoRep1000Half[[i]], NmpzHalf[i,], ThPathsRep1000Half[[i]])
}
```

```
# Calculate sd of DeltaTheta(j) for each position per replication
sdDThjRep1000Half <- lapply(ThPathsRep1000Half, function(ThPaths){apply(ThPaths$DThj, 1, sd)})
```

#### \(K=1000, N \sim 1130\)

```
# Define the dataset 
NmpzDouble <- floor(Nmpz*2)
num_SNPs <- nrow(NmpzDouble)
rownames(NmpzDouble) <- rownames(Nmpz)
NmpzDouble
```

```
# Generate microstates dataset
K <- 1000
GenoRep1000Double <- vector("list", num_SNPs)

for (i in 1:num_SNPs){
  set.seed(i)
  GenoRep1000Double[[i]] <- data.frame(replicate(K, sample(rep(c(-1,+1,0), times = c(NmpzDouble$Nm[i],NmpzDouble$Np[i],NmpzDouble$Nz[i]))), simplify = FALSE))
  colnames(GenoRep1000Double[[i]]) <- paste0("Rep",c(1:K))
}

# Call Fun_CumDistr
CumDistrRep1000Double <- vector("list", num_SNPs)
for (i in 1:num_SNPs) {
  CumDistrRep1000Double[[i]] <- Fun_CumDistr(GenoRep1000Double[[i]], NmpzDouble[i,])
}

# Call Fun_ThPaths
ThPathsRep1000Double <- vector("list", num_SNPs)
for (i in 1:num_SNPs){
  ThPathsRep1000Double[[i]] <- Fun_ThPaths(CumDistrRep1000Double[[i]])
}

# Call Fun_pGIFT
pGIFTRep1000Double <- vector("list", num_SNPs)
for (i in 1:num_SNPs){
  pGIFTRep1000Double[[i]] <- Fun_pGIFT(GenoRep1000Double[[i]], NmpzDouble[i,], ThPathsRep1000Double[[i]])
}
```

```
# Calculate sd of DeltaTheta(j) for each position per replication
sdDThjRep1000Double <- lapply(ThPathsRep1000Double, function(ThPaths){apply(ThPaths$DThj, 1, sd)})
```

### 3. Results - Generate plots

#### Figure 3A

Plot the null hypothesis, \(\Theta\)
paths, of all replications per SNP for \(N
\sim 565\), \(N \sim 280\),
\(N \sim 1130\). The black line
corresponds to the average value calculated per position.

```
# Define function to plot Theta paths
Fun_PlotThetaPaths <- function(ThPathsRep){
  library(ggplot2)
  num_SNPs <- length(ThPathsRep)
  
  j <- 1
  ThPaths <- ThPathsRep[[j]]
  # Plot the two paths: thetaj 
   
  Thj <- ThPaths$Thj
  Th0j <- ThPaths$Th0j
  DThj <- ThPaths$DThj
  N <- nrow(Thj)
  num_Reps <- ncol(Thj)
  
  j <- 1
  i <- 1
  # Plot the Thetaj path
  plot(c(1:N), Thj[,i], xlab = "j", ylab = "Theta-paths",
         xlim = c(1, nrow(Thj)), ylim = c(floor(min(Thj,na.rm = TRUE)),ceiling(max(Thj,na.rm = TRUE))),
         pch = 1, cex = 0.001, lwd =2, cex.lab = 1.6, cex.axis = 2, col = alpha("grey60", 0.2))
  for (i in 2:num_Reps) {
    points(c(1:N), Thj[,i], xlab = "j", ylab = "Theta-paths",
         xlim = c(1, nrow(Thj)), ylim = c(floor(min(Thj,na.rm = TRUE)),ceiling(max(Thj,na.rm = TRUE))),
         pch = 1, cex = 0.001, lwd =2, cex.lab = 1.6, cex.axis = 2,col = alpha("grey60", 0.2))
  }
  
  for (j in 2:num_SNPs) {
    ThPaths <- ThPathsRep[[j]]
    # Plot the two paths: thetaj 
     
    Thj <- ThPaths$Thj
    Th0j <- ThPaths$Th0j
    DThj <- ThPaths$DThj
    N <- nrow(Thj)
    num_Reps <- ncol(Thj)

    # Plot the Thetaj path
    for (i in 1:num_Reps) {
      points(c(1:N), Thj[,i], xlab = "j", ylab = "Theta-paths",
           xlim = c(1, nrow(Thj)), ylim = c(floor(min(Thj,na.rm = TRUE)),ceiling(max(Thj,na.rm = TRUE))),
           pch = 1, cex = 0.001, lwd =2, cex.lab = 1.6, cex.axis = 2, col = alpha("grey60", 0.2))
    }
  }
  
  for (j in 1:num_SNPs) {
    Thj <- ThPathsRep[[j]]$Thj
    points(c(1:N), apply(Thj,1,mean),pch = 1, cex = 0.001, lwd =2, cex.lab = 1.6, cex.axis = 2,col = "black")
  }
}

# Call the Fun_PlotThetaPaths for each sample size
par(mfrow=c(1,3))
Fun_PlotThetaPaths(ThPathsRep1000)
Fun_PlotThetaPaths(ThPathsRep1000Half)
Fun_PlotThetaPaths(ThPathsRep1000Double)
```

#### Figure 3B

Plot the null hypothesis, \(\Delta
\Theta\) paths, of all replications per SNP for \(N \sim 565\).

```
# Define function to plot Delta-Theta paths
Fun_PlotDeltaThetaPaths <- function(ThPaths, ymin, ymax, title){
  
  # Plot the two paths: thetaj 
  DThj <- ThPaths$DThj
  N <- nrow(DThj)
  num_SNPs <- ncol(DThj)
  
  i <- 1
  # Plot the difference between the two paths: DThj = Thetaj-Theta0j
  plot(seq(1, N, by = 1), DThj[,i], xlab = "j", ylab = "DThj", pch = 1, cex = 0.3, cex.lab = 1.8, cex.axis = 1.8, cex.main = 2,
       ylim = c(floor(ymin),ceiling(ymax)), main = title)
  for (i in 2:num_SNPs) {
    points(seq(1, N, by = 1), DThj[,i], xlab = "j", ylab = "DThj", pch = 1, cex = 0.3)
  }
}
```

```
# Set the limits on y-axis
ylim_min <- unlist(lapply(ThPathsRep1000, function(ThPaths){min(unlist(ThPaths$DThj), na.rm = TRUE)}))
ylim_max <- unlist(lapply(ThPathsRep1000, function(ThPaths){max(unlist(ThPaths$DThj), na.rm = TRUE)}))

# Call the Fun_PlotDeltaThetaPaths for sample size N~565
par(mfrow=c(1,6))
for (i in 1:num_SNPs) {
  Fun_PlotDeltaThetaPaths(ThPathsRep1000[[i]], min(ylim_min), max(ylim_max), paste("SNP",i))
}
```

#### Figure 3C

**Standard Deviation \(\Delta
\Theta(j) / \sqrt N\) against j/N**

```
# Plot the Standard Deviation $\Delta \Theta(j) / \sqrt N$ against j/N for all SNPs colored by the sample size
par(mfrow=c(1,6))
for (i in 1:num_SNPs) {
  N <- nrow(ThPathsRep1000[[i]]$DThj)
  plot(c(1:N)/N, sdDThjRep1000[[i]]/sqrt(N), pch = 1, cex = 0.5, cex.lab = 2, cex.axis = 2, cex.main = 2, col = "black", 
       main = rownames(Nmpz)[i], xlab = "j/N", ylab = "sd DThj/sqrt(N)",
       ylim = c(min(c(unlist(sdDThjRep1000),unlist(sdDThjRep1000Half),unlist(sdDThjRep1000Double)), na.rm = TRUE),
                max(c(unlist(sdDThjRep1000),unlist(sdDThjRep1000Half),unlist(sdDThjRep1000Double)), na.rm = TRUE))/sqrt(N))
  N <- nrow(ThPathsRep1000Half[[i]]$DThj)
  points(c(1:N)/N, sdDThjRep1000Half[[i]]/sqrt(N), pch = 1, cex = 0.5, col = "red3")
  N <- nrow(ThPathsRep1000Double[[i]]$DThj)
  points(c(1:N)/N, sdDThjRep1000Double[[i]]/sqrt(N), pch = 1, cex = 0.5, col = "blue3")
  legend("topright", legend = c("N~565","N~280","N~1130"), fill = c("black","red3","blue3"), cex = 1.4)
}
```

### Figure 6C

**Boxplots of ranges of \(-log\_{10}\hat{p}GIFT\)**

For each sample size, we set a threshold based on the \(99\%\) and \(95\%\) percentiles of the range of values
across replications per SNP. Therefore, we calculate the average of the
values across the six SNPs.

```
# Define function to plot the boxplots of -log10pGIFT
Fun_pGIFTBoxplot <- function(pGIFT, xlab, ylim, title){
  threshold099 <- lapply(pGIFT, function(x){quantile(-log10(x), 0.99)})
  threshold095 <- lapply(pGIFT, function(x){quantile(-log10(x), 0.95)})
  boxplot(lapply(pGIFT, function(x){-log10(x)}), ylab = "-log10(pGIFT)", xaxt = "n", 
          ylim = c(floor(min(0,ylim[1])),ceiling(ylim[2])), main = title, cex.lab = 2, cex.axis = 2)
  text(c(1:length(pGIFT)), par("usr")[3] - 3, xlab, srt = 0, cex = 2, xpd = TRUE)
  abline(h=mean(unlist(threshold099)), col = "red", lty = "dashed", lwd = 2)
  abline(h=mean(unlist(threshold095)), col = "red", lty = "dashed", lwd = 2)
}
```

```
# Set the limit on y-axis
y_bp_pGIFT_min_max <- c(-log10(max(c(unlist(pGIFTRep1000),unlist(pGIFTRep1000Half),unlist(pGIFTRep1000Double)))),
                        -log10(min(c(unlist(pGIFTRep1000),unlist(pGIFTRep1000Half),unlist(pGIFTRep1000Double)))))

# Call the Fun_pGIFTBoxplot for each sample size
par(mfrow=c(1,3))
Fun_pGIFTBoxplot(pGIFTRep1000, rownames(Nmpz), y_bp_pGIFT_min_max, "K=1000 Replications, N~565")
Fun_pGIFTBoxplot(pGIFTRep1000Half, rownames(NmpzHalf), y_bp_pGIFT_min_max, "K=1000 Replications, N~280")
Fun_pGIFTBoxplot(pGIFTRep1000Double, rownames(NmpzDouble), y_bp_pGIFT_min_max, "K=1000 Replications, N~1130")
```

### Figure 6D

**Boxplots of ranges of \(-log\_{10}pGIFT\)**

For each sample size, we set a threshold based on the \(99\%\) and \(95\%\) percentiles of the range of values
across replications per SNP. Therefore, we calculate the average of the
values across the six SNPs.

```
# Define function to plot the boxplots of -log10pGIFT
Fun_pSNPadjBHBoxplot <- function(pGIFT, pGIFTadjBH, xlab, ylim, title){
  threshold099 <- lapply(pGIFT, function(x){quantile(-log10(x), 0.99)})
  threshold095 <- lapply(pGIFT, function(x){quantile(-log10(x), 0.95)})
  boxplot(lapply(pGIFTadjBH, function(x){-log10(x)}), ylab = "-log10(pGIFT)", xaxt = "n", 
          ylim = c(floor(min(0,ylim[1])),ceiling(ylim[2])), main = title, cex.lab = 2, cex.axis = 2)
  text(c(1:length(pGIFT)), par("usr")[3] - 3, xlab, srt = 0, cex = 2, xpd = TRUE)
  abline(h=mean(unlist(threshold099)), col = "red", lty = "dashed", lwd = 2)
  abline(h=mean(unlist(threshold095)), col = "red", lty = "dashed", lwd = 2)
}
```

```
# Set the limit on y-axis
y_bp_pGIFT_min_max <- c(-log10(max(c(unlist(pGIFTRep1000),unlist(pGIFTRep1000Half),unlist(pGIFTRep1000Double)))),
                        -log10(min(c(unlist(pGIFTRep1000),unlist(pGIFTRep1000Half),unlist(pGIFTRep1000Double)))))

# Call the Fun_pSNPadjBHBoxplot for each sample size
par(mfrow=c(1,3))
Fun_pSNPadjBHBoxplot(pGIFTRep1000, lapply(pGIFTRep1000, function(x){p.adjust(x, method = "BH")}), rownames(Nmpz), y_bp_pGIFT_min_max, "K=1000 Replications, N~565")
Fun_pSNPadjBHBoxplot(pGIFTRep1000Half, lapply(pGIFTRep1000Half, function(x){p.adjust(x, method = "BH")}), rownames(NmpzHalf), y_bp_pGIFT_min_max, "K=1000 Replications, N~280")
Fun_pSNPadjBHBoxplot(pGIFTRep1000Double, lapply(pGIFTRep1000Double, function(x){p.adjust(x, method = "BH")}), rownames(NmpzDouble), y_bp_pGIFT_min_max, "K=1000 Replications, N~1130")
```
