## Supplementary material for "Investigative power of Genomic Informational Field Theory (GIFT) relative to GWAS for genotype-phenotype mapping": S5

### Slide 1
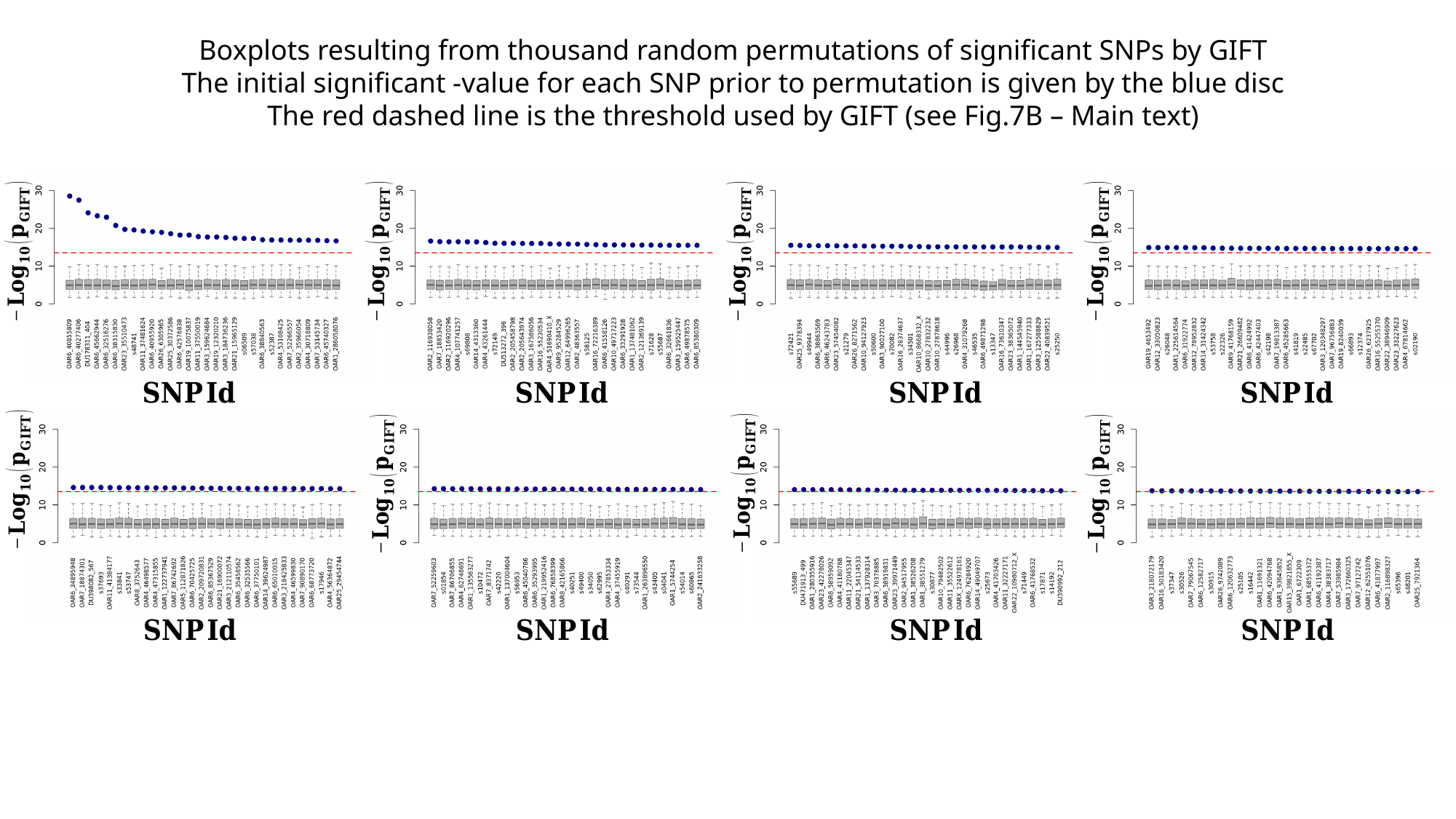

### Slide 2
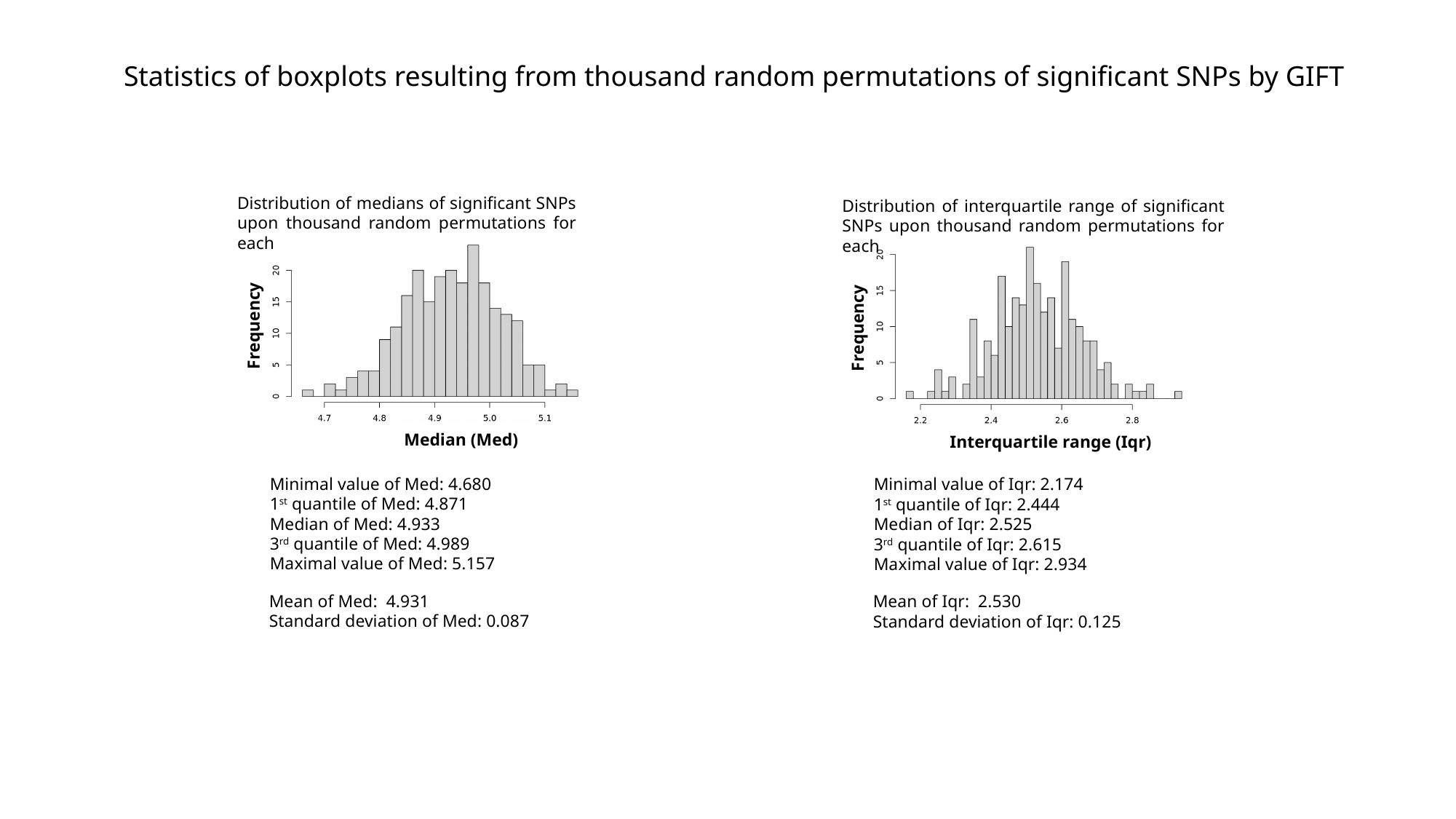

Statistics of boxplots resulting from thousand random permutations of significant SNPs by GIFT
Distribution of medians of significant SNPs upon thousand random permutations for each
Distribution of interquartile range of significant SNPs upon thousand random permutations for each
Frequency
Frequency
Median (Med)
Interquartile range (Iqr)
Minimal value of Med: 4.680
1^st^ quantile of Med: 4.871
Median of Med: 4.933
3^rd^ quantile of Med: 4.989
Maximal value of Med: 5.157
Minimal value of Iqr: 2.174
1^st^ quantile of Iqr: 2.444
Median of Iqr: 2.525
3^rd^ quantile of Iqr: 2.615
Maximal value of Iqr: 2.934
Mean of Med: 4.931
Standard deviation of Med: 0.087
Mean of Iqr: 2.530
Standard deviation of Iqr: 0.125
