## Supplementary material for "Investigative power of Genomic Informational Field Theory (GIFT) relative to GWAS for genotype-phenotype mapping": S7

Scottish Blackface Sheep Analysis: Run GIFT with residuals of model using genotype coming from filters by GWAS


### Scottish Blackface Sheep Analysis: Run GIFT with residuals of model using genotype coming from filters by GWAS

###### Supplemental material by Kyratzi et al.

```
#### CODE to run in terminal ####

# set the working directory
cd set/your/path 
  
# Load PLINK
module load plink

# Filter the dataset
plink --file Sheep --noweb --allow-no-sex --autosome-num 26 --make-bed --out Sheep
plink --noweb --file Sheep  --geno 0.10  --hwe 0.000001  --maf 0.05  --autosome-num 26 --allow-no-sex --make-bed --out Sheep05_724
```

### Perform GWAS analysis

```
# Perform GWAS analysis

# Calculate the relationship matrix of the individuals
./gemma-0.98.5 -bfile Sheep05_724 -maf 0 -miss 0.1 -gk 1 -o Rmatrix

# The residuals of the natural log-transformed metabolites SAM, mB12, aB12, DMG, TMG, MMA, PPA are on the 7th to 13th column of the .ped file
./gemma-0.98.5 -bfile Sheep05_724 -n 3 -hwe 0 -maf 0 -miss 1 -k Rmatrix.cXX.txt -lmm 4 -o GWAS_rbon_area_05

# Export GWAS results
# Export columns of chromosome, name, position and pwald
cd output
cat GWAS_rbon_area_05.assoc.txt | awk '{print $1, $2, $3, $13}' > GWAS_rbon_area_05.assoc_subset.txt
```

```
# Calculate the number of independent SNPs in the dataset

module load plink
plink -file Sheep05_724 --noweb --allow-no-sex --autosome-num 26 --recode vcf-iid --out data
# --noweb: This flag disables web-based operations.
# --allow-no-sex: This flag allows samples with missing sex information.
# --autosome-num 26: This sets the number of autosomes in the dataset to 26. The autosomes are the non-sex chromosomes.
# --recode vcf-iid: This option specifies the output format. In this case, it's requesting the data to be recoded in the VCF (Variant Call Format) with individual identifiers.
# --out data: This specifies the prefix for the output files. 

module load bcftools
bcftools +prune --max r2=0.1 --window 1Mb -Ov -o testoutput.vcf data.vcf
# +prune: This option is used to perform LD (linkage disequilibrium) pruning on the input VCF file. LD pruning is a common step in population genetics to reduce redundancy in genetic data by removing correlated variants.
# --max r2=0.1: This sets the maximum r-squared (a measure of correlation between genetic variants) allowed for variants to be considered correlated. Variants with an r-squared above 0.1 will be pruned (removed).
# --window 1Mb: This defines the size of the sliding window for LD pruning. Variants within this window will be considered for LD pruning. In this case, the window size is set to 1 megabase (1Mb).
# -Ov: This option specifies the output format as VCF.
# -o testoutput.vcf: This specifies the output file name. The pruned VCF data will be saved to a file named testoutput.vcf.
# data.vcf: This is the input VCF file on which LD pruning will be performed.

bcftools stats testoutput.vcf > testoutput.vcf.stats
# stats: The subcommand that computes various statistics for a VCF file.
# testoutput.vcf: The input VCF file for which statistics are being computed.
```

### Perform GIFT analysis

```
# set the working directory
# setwd(YourPath)

require(data.table) #for fread
Sheep05_724 <- fread("Sheep05_724.ped", header = FALSE)
Sheep05_724.map <- fread("Sheep05_724.map", header = FALSE)
```

```
# Read the pheno dataset
CT_traits_724_pc_res <- read.csv("CT_traits_724_pc_res.csv")

# match the individuals in the the two datasets
CT_traits_724_pc_res <- CT_traits_724_pc_res[match(unlist(Sheep05_724.fam[,2]),CT_traits_724_pc_res$id),]
```

```
# replace the values with the microstates -1, +1, 0
# GG=CC=-1
# TT=AA=+1
# AG=CG=AC=AT=0
# unique(as.vector(unlist(genotype)))

library(pegas)
genotype <- alleles2loci(Sheep05_724[,-c(1:6)])

library(dplyr)
Geno <- as.data.frame(matrix(case_when((genotype=="G/G") | (genotype=="C/C") ~ -1,
                                       (genotype=="T/T") | (genotype=="A/A") ~ +1,
                                       genotype=="0/0" ~ NA_real_,
                                       TRUE ~ 0),
                             nrow = nrow(genotype), ncol = ncol(genotype), byrow = FALSE))

# Set the row and column names of the new geno dataset
rownames(Geno) <- unlist(Sheep05_724[,2])
colnames(Geno) <- unlist(Sheep05_724.map[,2])
```

#### Treat missing phenotypic values

```
# Function to remove the individuals with missing values on the phenotype (residuluals) from phenotype list and genotype dataset
# Input variables:
#   Phenotype: dataframe consisting of info about individuals and the value of phenotype
#   Genotype: dataframe consisting of microstates (row: individuals, col: SNPs)
# Output variables:
#   List of phenotype and genotype datasets (row: individuals, col: phenotype) after removing individuals with missing phenotype

Fun_filterMissingPheno <- function(Phenotype, Genotype, namePheno){
  ind.missPheno <- which(is.na(Phenotype[,namePheno]))
  Phenotype <- Phenotype[-ind.missPheno,c("id", namePheno)]
  Genotype <- Genotype[-ind.missPheno,]
  
  return(list(Pheno = Phenotype, Geno = Genotype))
}

# Call Fun_filterMissingPheno
# Data_noMissPheno <- Fun_filterMissingPheno(CT_traits_724_pc_res, Geno, "rbon_area")
```

```
# Function to calculate the number of each microstate for all SNPs
# Input variables:
#   Geno: genotype dataset (row: individuals, col: SNPs)
# Output variables:
#   Nmpz: dataframe of number of microstates for each SNP

Function_Nmpz <- function(Geno){

  Nm <- data.frame(Nm = apply(Geno, 2, function(x){sum(x==-1, na.rm = TRUE)}))
  Np <- data.frame(Np = apply(Geno, 2, function(x){sum(x==1, na.rm = TRUE)})) 
  Nz <- data.frame(Nz = apply(Geno, 2, function(x){sum(x==0, na.rm = TRUE)}))
  Nmpz <- cbind(Nm,Np,Nz)
  
  return(Nmpz)
}

# Call Function_Nmpz
Nmpz <- Function_Nmpz(Data_noMissPheno$Geno)
```

#### Pre-processing of the datasets to run GIFT analysis

Order the individuals according to their phenotype and create the
sorted datasets.

```
# Function to order the individuals according to their phenotype and create the sorted datasets
# Input variables:
#   Geno: geno dataset (row: individuals, col: SNPs)
#   Pheno: pheno dataset (row: individuals, col: id, phenotype)
# Output variables:
#   List of sorted geno (row: individuals, col: SNPs) and pheno datasets (row: individuals, col: phenotype)

Fun_sortData <- function(Geno, Pheno){
  GenoSorted <- data.frame(matrix(NA, ncol = ncol(Geno), nrow = nrow(Pheno)))
  PhenoSorted <- data.frame(matrix(NA, ncol = ncol(Geno), nrow = nrow(Pheno)))
  IDSorted <- data.frame(matrix(NA, ncol = ncol(Geno), nrow = nrow(Pheno)))
  for (i in 1:ncol(Geno)) {
    no.NA <- !is.na(Geno[,i]) # identify the non-missing values in the gene list for the SNP i
    pheno.ordered <- sort(Pheno[no.NA,2], decreasing = FALSE, index.return = TRUE)
    GenoSorted[c(1:sum(no.NA)),i] <- (Geno[no.NA,i])[pheno.ordered$ix]
    PhenoSorted[c(1:sum(no.NA)),i] <- pheno.ordered$x
    IDSorted[c(1:sum(no.NA)),i] <- (Pheno[no.NA,1])[pheno.ordered$ix]
    
    colnames(GenoSorted) <- colnames(Geno)
    colnames(PhenoSorted) <- colnames(Geno)
    colnames(IDSorted) <- colnames(Geno)
  }
  
  return(list(Geno = GenoSorted, Pheno = PhenoSorted, id = IDSorted))
}

# Call Fun_sortData
sortData <- Fun_sortData(Data_noMissPheno$Geno, Data_noMissPheno$Pheno)
```

#### GIFT analysis computations

Compute the cumulative distributions of the ordered and random
configuration to formulate the \(\Theta\)-paths.

```
# Function to count the number of individuals in each bin for each SNP
# Input variables:
#   Geno: microstate dataset (row: individuals, col: SNPs)
#   Pheno: sorted pheno dataset (row: individuals, col: phenotype)
# Output variables:
#   CountValBins: dataframe with the number of how many individuals in each bin (rows) per SNP (columns)
# Function to count the number of individuals in each bin for each gene

Fun_CountValBins <- function(Geno, Pheno){
  
  CountValBins <- Map(function(Geno, Pheno){
    Geno <- unlist(Geno)
    Pheno <- unlist(Pheno)
    
    CountValBins <- rep(NA, length(Geno))
    no.NA <- !is.na(Geno)
    N <- sum(no.NA)
    phen <- Pheno[no.NA]
    
    numBins <- 1 
    k <- 1
    
    CountValBins[1:length(as.vector(table(phen)))] <- as.vector(table(phen))
    
    return(CountValBins)}, as.list(Geno), as.list(Pheno))
  
  CountValBins <- do.call(cbind, lapply(CountValBins, function(x){matrix(x, nrow = nrow(Geno), byrow = FALSE)}))
  
  rownames(CountValBins) <- paste("Bin", c(1:nrow(Geno)))
  colnames(CountValBins) <- colnames(Geno)
  
  return(as.data.frame(CountValBins))
}

# Call Fun_CountValBins
CountValBins <- Fun_CountValBins(sortData$Geno, sortData$Pheno)
```

```
# Function to compute the cumulative distributions of the ordered and random configuration according to the bins for each SNP
# Input variables:
#   Geno: sorted microstate dataset (row: individuals, col: SNPs)
#   CountValBins:  dataframe with the number of how many individuals in each bin (rows) per SNP (columns)
#   Nmpz: dataframe of number of microstates for each SNP (rows)
# Output variables:
#   List of Wp, Wm, Wz, W0p, W0m, W0z: Cumulative distributions of +1,-1 respectively for ordered and random configuration according to the bins for each SNP

Fun_CumDistr <- function(Geno, CountValBins, Nmpz){
  library(purrr) # for accumulate
  
  CumDistrW <- Map(function(Geno,CountValBins){
    bins <- CountValBins[!is.na(CountValBins)]
    numBins <- length(bins)
    
    # Split the vector of microstates into groups per bin
    grouped_geno <- split(Geno, rep(seq_along(bins), bins))
    # Remove NA values from each vector in the list
    grouped_geno <- lapply(grouped_geno, function(x) na.omit(x))
    
    # Define a function to calculate the weighted cumulative sums
    calculate_weighted_cumulative_sums_per_bin <- function(grouped_geno) {
      weights <- (1:length(grouped_geno)) / length(grouped_geno)
      group_cumsum <- sapply(-1:1, function(group) {
        weights * sum(grouped_geno == group)
      })
      return(group_cumsum)
    }
    
    # Calculate the weighted cumulative sums for each group per bin
    cumulative_sums <- t(matrix(sapply(grouped_geno, calculate_weighted_cumulative_sums_per_bin)))
    cumulative_sums <- do.call(rbind, lapply(cumulative_sums, function(x){matrix(x, ncol = 3, byrow = FALSE)}))
    
    # Add the cumulative sums from previous bins
    result <- apply(cumulative_sums, 2, function(x){
      accumulate(2:numBins, function(x, n) {
        indstart <- sum(bins[1:(n-1)]) + 1
        indend <- sum(bins[1:n])
        s <- x[indstart - 1]
        x[indstart:indend] <- x[indstart:indend] + s
        return(x)}, .init = x)})
    
    Wm <- result[[1]][[length(result[[1]])]]
    Wz <- result[[2]][[length(result[[2]])]]
    Wp <- result[[3]][[length(result[[3]])]]
    
    return(data.frame(Wp = Wp, Wm = Wm, Wz = Wz))
    
  }, Geno, CountValBins)
  
  Wp <- as.data.frame(do.call(qpcR:::cbind.na, lapply(CumDistrW, `[[`, "Wp")))
  Wm <- as.data.frame(do.call(qpcR:::cbind.na, lapply(CumDistrW, `[[`, "Wm")))
  Wz <- as.data.frame(do.call(qpcR:::cbind.na, lapply(CumDistrW, `[[`, "Wz")))
  
  CumDistrW0 <- apply(Nmpz, 1, function(Nmpz){
    Nmpz <- unlist(Nmpz)
    N <- sum(Nmpz)
    W0p <- c(1:N) * Nmpz[2] / N
    W0m <- c(1:N) * Nmpz[1] / N
    W0z <- c(1:N) * Nmpz[3] / N
    return(data.frame(W0p = W0p, W0m = W0m, W0z = W0z))
  })
  
  W0p <- as.data.frame(do.call(qpcR:::cbind.na, lapply(CumDistrW0, `[[`, "W0p")))
  W0m <- as.data.frame(do.call(qpcR:::cbind.na, lapply(CumDistrW0, `[[`, "W0m")))
  W0z <- as.data.frame(do.call(qpcR:::cbind.na, lapply(CumDistrW0, `[[`, "W0z")))
  
  return(list(Wp = Wp, Wm = Wm, Wz = Wz, W0p = W0p, W0m = W0m, W0z = W0z))
}


# Call Fun_CumDistr
CumDistr <- Fun_CumDistr(sortData$Geno, CountValBins, Nmpz)
```

#### pGIFT for two and three states

For +1,-1:

\[pGIFT =
\Bigg(\frac{1}{2}-\frac{1}{\pi}tan^{-1} \Big(\frac{\sqrt{-2\phi
\tilde\phi}}{2(\phi-\tilde\phi)} \Big) \Bigg) \exp\Big\{
-\frac{(\phi-\tilde\phi)^2}{2Nw(1-w)} \Big\}, w=N\_+/N, \quad \phi =
\max\_{j}\Delta\Theta(j) , \quad \tilde\phi = \min\_{j}\Delta\Theta(j)
\]

\[=
\Bigg(\frac{1}{2}-\frac{1}{\pi}tan^{-1} \Big(\frac{\sqrt{-2\phi
\tilde\phi}}{2(\phi-\tilde\phi)} \Big) \Bigg) \exp\Big\{
-\frac{N(\phi-\tilde\phi)^2}{2N\_+N\_-} \Big\} \]

For +1,0:

\[pGIFT =
\Bigg(\frac{1}{2}-\frac{1}{\pi}tan^{-1} \Big(\frac{\sqrt{-2\phi
\tilde\phi}}{2(\phi-\tilde\phi)} \Big) \Bigg) \exp\Big\{
-\frac{2(\phi-\tilde\phi)^2}{Nw(1-w)} \Big\}, w=N\_+/N, \quad \phi =
\max\_{j}\Delta\Theta(j) , \quad \tilde\phi = \min\_{j}\Delta\Theta(j)
\]

\[=
\Bigg(\frac{1}{2}-\frac{1}{\pi}tan^{-1} \Big(\frac{\sqrt{-2\phi
\tilde\phi}}{2(\phi-\tilde\phi)} \Big) \Bigg) \exp\Big\{
-\frac{2N(\phi-\tilde\phi)^2}{N\_+N\_0} \Big\} \]

For -1,0:

\[pGIFT =
\Bigg(\frac{1}{2}-\frac{1}{\pi}tan^{-1} \Big(\frac{\sqrt{-2\phi
\tilde\phi}}{2(\phi-\tilde\phi)} \Big) \Bigg) \exp\Big\{
-\frac{2(\phi-\tilde\phi)^2}{Nw(1-w)} \Big\}, w=N\_-/N, \quad \phi =
\max\_{j}\Delta\Theta(j) , \quad \tilde\phi =
\min\_{j}\Delta\Theta(j)\] \[ =
\Bigg(\frac{1}{2}-\frac{1}{\pi}tan^{-1} \Big(\frac{\sqrt{-2\phi
\tilde\phi}}{2(\phi-\tilde\phi)} \Big) \Bigg) \exp\Big\{
-\frac{2N(\phi-\tilde\phi)^2}{N\_-N\_0} \Big\}\]

For -1,+1,0:

\[pGIFT =
\Bigg(\frac{1}{2}-\frac{1}{\pi}tan^{-1} \Big(\frac{\sqrt{-\phi
\tilde\phi}}{(\phi-\tilde\phi) \sqrt{2}} \Big) \Bigg) \exp\Big\{
-\frac{8N(\phi-\tilde\phi)^2}{N(N-N\_0)-(N\_+-N\_-)^2} \Big\}, \quad \phi =
\max\_{j}\Delta\Theta(j) , \quad \tilde\phi =
\min\_{j}\Delta\Theta(j)\]

```
# Function to calculate the p-value
# Input variables:
#   Geno: sorted geno dataset (row: individuals, col: SNPs)
#   Nmpz: dataframe of number of microstates for each SNP
#   ThPaths: List of Theta-paths (output of Fun_ThPaths)
# Output variables:
#   pGIFT: vector consists of pGIFT values

Fun_pGIFT <- function(Geno, Nmpz, ThPaths){
  pGIFT <- c()
  
  num_SNPs <- ncol(Geno)
  DThj <- as.data.frame(ThPaths$DThj)
  
  phi <- apply(DThj, 2, function(x){max(x, na.rm = TRUE)})
  phi_tilde <- apply(DThj, 2, function(x){min(x, na.rm = TRUE)})
  
  for (i in 1:num_SNPs) {
    
    no.NA <- !is.na(Geno[,i])
    N <- sum(no.NA)
    
    if (sum(Nmpz[i,]>=3)==3) { #check if we have three states - compute the pGIFT for three-states
      e <- exp( -(8*N*(phi[i]-phi_tilde[i])^2) / (N*(N-Nmpz[i,3])-(Nmpz[i,2]-Nmpz[i,1])^2) )
      
      pGIFT[i] <- ( (1/2)-(1/pi)*atan(sqrt(-phi[i]*phi_tilde[i])/((phi[i]-phi_tilde[i])*sqrt(2))) ) *e
      
    } #end if condition for three states
    if (sum(Nmpz[i,]>=3)==2) {  # length(uniq.states)==2
      uniq.states <- data.frame(table(Geno[no.NA,i]))
      ind_geno <- which(uniq.states$Freq>=3)
      if (length(intersect(c(-1,0),uniq.states[ind_geno,1]))==2){ #check if we have -1 and 0 states to use Thjm and Thjz
        e <- exp( -(2*N*(phi[i]-phi_tilde[i])^2) / (uniq.states$Freq[ind_geno[1]]*uniq.states$Freq[ind_geno[2]]) )
        
        pGIFT[i] <- ( (1/2)-(1/pi)*atan(sqrt(-2*phi[i]*phi_tilde[i])/(2*(phi[i]-phi_tilde[i]))) ) *e
      } #end if condition for two states: -1, 0
      
      if (length(intersect(c(+1,0),uniq.states[ind_geno,1]))==2){ #check if we have +1 and 0 states to use Thjp and Thjz
        e <- exp( -(2*N*(phi[i]-phi_tilde[i])^2) / (uniq.states$Freq[ind_geno[1]]*uniq.states$Freq[ind_geno[2]]) )
        
        pGIFT[i] <- ( (1/2)-(1/pi)*atan(sqrt(-2*phi[i]*phi_tilde[i])/(2*(phi[i]-phi_tilde[i]))) ) *e
      } #end if condition for two states: +1, 0
      
      else if (length(intersect(c(-1,+1),uniq.states[ind_geno,1]))==2){ #check if we have -1 and +1 states to use Thjm and Thjp
        e <- exp( -(N*(phi[i]-phi_tilde[i])^2) / (2*uniq.states$Freq[ind_geno[1]]*uniq.states$Freq[ind_geno[2]]) )
        
        pGIFT[i] <- ( (1/2)-(1/pi)*atan(sqrt(-2*phi[i]*phi_tilde[i])/(2*(phi[i]-phi_tilde[i]))) ) *e
      } #end if condition for two states: -1, +1
    } #end if condition for two states
    
    if (sum(Nmpz[i,]>=3)<2) { #check if we have only one state
      pGIFT[i] <- 1 # set this to one so after taking the logarithm we have zero (non-significant)
    }
    
  }
  
  return(pGIFT)
  
}

# Call Fun_pGIFT
pGIFT <- Fun_pGIFT(sortData$Geno, Nmpz, ThPaths)
```

### RESULTS

#### Figure 4

```
require(data.table)
MapResults <- fread("MapResults_rbon_area.csv")
ThPaths <- readRDS("ThPaths_rbon_area.rds")
sortData <- readRDS("sortData_rbon_area.rds")
Nmpz <- read.csv("Nmpz_rbon_area.csv", row.names = 1)
```

```
# Function to generate plots of theta paths
Fun_PlotPaths <- function(sortData, Nmpz, ThPaths, MapResults, indToPlot){
  
  # Plot the two paths: thetaj and theta0j
  set.seed(1)
  
  Thj <- ThPaths$Thj
  Th0j <- ThPaths$Th0j
  DThj <- ThPaths$DThj
  
  Geno <- sortData$Geno
  N <- nrow(Geno)
  i <- 0
  # par(mfrow = c(1,2))
  for (n in indToPlot) {
    i <- i+1
    # Plot the two paths: thetaj and theta0j
    ind.p <- which(Geno[,n]==1)
    ind.z <- which(Geno[,n]==0)
    ind.m <- which(Geno[,n]==-1)
    randomgenes <- sample(Geno[,n], replace = FALSE)
    indr.p <- which(randomgenes==1)
    indr.z <- which(randomgenes==0)
    indr.m <- which(randomgenes==-1)
    plot(ind.p, Thj[ind.p,n], xlab = "i", ylab = "Theta-paths", main = paste0("SNP",i),
         xlim = c(1, nrow(Thj)), ylim = c(floor(min(Thj[,n],na.rm = TRUE)),ceiling(max(Thj[,n],na.rm = TRUE))),
         pch = 1, cex = 0.8, lwd =2, col="firebrick1", cex.lab = 1.6, cex.axis = 2, cex.main = 2)
    points(ind.m, Thj[ind.m,n], pch = 1, cex = 0.8, lwd = 2, col="deepskyblue4")
    points(ind.z, Thj[ind.z,n], pch = 1, cex = 0.8, lwd = 2, col="gray30")
    points(indr.p, Th0j[indr.p,n], pch = 1, cex = 0.8, lwd = 2, col="firebrick1")
    points(indr.m, Th0j[indr.m,n], pch = 1, cex = 0.8, lwd = 2, col="deepskyblue4")
    points(indr.z, Th0j[indr.z,n], pch = 1, cex = 0.8, lwd = 2, col="gray30")
    # legend("left", legend = c("+1 states", "-1 states", "0 states"), fill = c("firebrick1", "deepskyblue4", "gray30"), cex = 1)
    abline(h=0, lty = 2, lwd = 1.5)
    
    # Plot the difference between the two paths: DThj = Thetaj-Theta0j
    plot(seq(1, N, by = 1), DThj[,n], xlab = "i", ylab = "DThj",  main = paste0("SNP",i),
         pch = 1, cex = 0.8, cex.lab = 1.6, cex.axis = 2, cex.main = 2,
         ylim = c(min(-sqrt(((N-c(1:N))*c(1:N))/N),DThj[,n],na.rm = TRUE), max(sqrt(((N-c(1:N))*c(1:N))/N),DThj[,n],na.rm = TRUE)))
    abline(h=0, lty = 2, lwd = 1.5)
  }
}
```

```
par(mfrow=c(3,4))
# Call Fun_PlotPaths
Fun_PlotPaths(sortData,Nmpz,ThPaths,MapResults,indSNPs)
```

#### Figure 5

```
# Function to generate plots of the effect size
Fun_plotEffectSize <- function(sortData, indToPlot){
  library(ggplot2)
  
  n <- 0
  for (i in indToPlot) {
    n <- n+1
    
    df <- data.frame(rbon_area = sortData$Pheno[,i], genotype = sortData$Geno[,i])
    # df$microstates <- factor(as.integer(df$genotype))
    df$microstates <- factor(df$genotype, levels = c("-1","0","1"), labels = c("-1","0","1"))

    lm <- lm(rbon_area~genotype, data = df) # create  the linear regression
    lmsumm <- summary(lm) #summary of the linear regression
    
    mypalette <- c("-1"="deepskyblue4", "0"="gray30", "1"="firebrick1")
    ggPlot <- ggplot(df, aes(x = genotype, y = rbon_area, col = microstates)) +
      geom_point(show.legend = FALSE, shape=1, size = 4) + xlim(-1,1) +
      #straight line using the intercept and the slope from the linear regression
      geom_abline(slope = lmsumm$coefficients[2,1], intercept = lmsumm$coefficients[1,1]) +
      scale_color_manual(values=mypalette) +
      theme_classic(base_size = 15)+
      ggtitle(paste0("SNP",n))
    print(ggPlot)
  }
}
```

```
# Call Fun_plotEffectSize
Fun_plotEffectSize(sortData,indSNPs)
```

#### Figure 7A & 7B

```
# Read GWAS results
pGWAS_rbone_area <- read.table("output/GWAS_rbon_area_05.assoc_subset.txt", header = TRUE)
pGWAS_rbone_area$mlog10_pGWAS <- -log10(pGWAS_rbone_area$p_wald)
```

```
# Read GIFT results
require(data.table)
MapResults_rbone_area <- fread("MapResults_rbon_area.csv")
```

```
# Read the thresholds values as calculated by the permutation analysis
threshold099 <-readRDS("threshold099N565.rds")
threshold095 <-readRDS("threshold095N565.rds")
```

```
library(qqman)
# Function to generate Manhattan plot for GWAS analyses
Fun_GWASManhattanPlots <- function(pGWAS_r, metabolite){
  num_IndepSNPs <- 10433 # set the number of independent SNPs as calculated using bcftools
  manhattan(pGWAS_r, chr = "chr", bp = "ps", p = "mlog10_pGWAS", snp = "rs",
            chrlabs = as.character(unique(pGWAS_r$chr)), logp = FALSE, ylab = "-log10(pGWAS)", 
            cex = 0.8, cex.axis = 2, cex.lab = 2, cex.main = 2, ylim = c(0,10),  col = c("blue3", "orange3"),
            main = paste0("GWAS_r", metabolite), suggestiveline = FALSE, genomewideline = FALSE)
  abline(h = -log10(0.01/num_IndepSNPs), col = "red", lty = "dashed", lwd = 2)
  abline(h = -log10(0.05/num_IndepSNPs), col = "red", lty = "dashed", lwd = 2)
}

# Function to generate Manhattan plot for GIFT analyses
Fun_GIFTManhattanPlots <- function(MapResults_r, metabolite, threshold1, threshold2){
manhattan(MapResults_r, chr = "CHRM", bp = "POSITION", p = "mlog10_pGIFT_adjBH", snp = "NAME",
            chrlabs = as.character(unique(MapResults_r$CHRM)), logp = FALSE, ylab = "-log10(pGIFT)", 
          cex = 0.8, cex.axis = 2, cex.lab = 2, cex.main = 2, ylim = c(0,30),  col = c("blue3", "orange3"),
            main = paste0("GIFT_r", metabolite), suggestiveline = FALSE, genomewideline = FALSE)
  abline(h = threshold1, col = "red", lty = "dashed", lwd = 2)
  abline(h = threshold2, col = "red", lty = "dashed", lwd = 2)
}

# Call Fun_GWASManhattanPlots and Fun_GIFTManhattanPlots
par(mfrow=c(1,2))

Fun_GWASManhattanPlots(pGWAS_rbone_area, "bone_area")
Fun_GIFTManhattanPlots(MapResults_rbone_area, "bone_area", threshold099, threshold095)
```

#### Figure 7C

```
num_IndepSNPs <- 10433 # set the number of independent SNPs as calculated using bcftools

MapResults_rbone_area_CHRM6 <- MapResults_rbone_area[MapResults_rbone_area$CHRM==6,]
highSNPsList <- list(GWAS = pGWAS_rbone_area$rs[pGWAS_rbone_area$mlog10_pGWAS>-log10(0.01/num_IndepSNPs)],
                     GIFT = MapResults_rbone_area_CHRM6$NAME[MapResults_rbone_area_CHRM6$mlog10_pGIFT_adjBH>threshold099])
```

#### Figure 7D

```
# Function to generate plots of theta paths
Fun_PlotDThPath <- function(sortData, Nmpz, ThPaths, MapResults, indToPlot){
  
  DThj <- ThPaths$DThj
  N <- nrow(DThj)
    
  # Plot the difference between the two paths: DThj = Thetaj-Theta0j
  plot(seq(1, N, by = 1), DThj[,indToPlot], xlab = "i", ylab = "DThj",  main = MapResults$NAME[indToPlot],
       pch = 1, cex = 0.8, cex.lab = 1.5, cex.axis = 1.8, cex.main = 1.8,
       ylim = c(min(-sqrt(((N-c(1:N))*c(1:N))/N),DThj[,indToPlot],na.rm = TRUE), max(sqrt(((N-c(1:N))*c(1:N))/N),DThj[,indToPlot],na.rm = TRUE)))
  abline(h=0, lty = 2, lwd = 1.5)
}

# Function to generate plots of the effect size
Fun_plotEffectSize <- function(sortData, MapResults, indToPlot){
  library(ggplot2)
    
  df <- data.frame(rbon_area = sortData$Pheno[,indToPlot], genotype = sortData$Geno[,indToPlot])
  # df$microstates <- factor(as.integer(df$genotype))
  df$microstates <- factor(df$genotype, levels = c("-1","0","1"), labels = c("-1","0","1"))
    
  mypalette <- c("-1"="deepskyblue4", "0"="gray30", "1"="firebrick1")
  ggPlot <- ggplot(df, aes(x = genotype, y = rbon_area, col = microstates)) +
    geom_point(show.legend = FALSE, shape=1, size = 4) + xlim(-1,1) +
    scale_color_manual(values=mypalette) +
    theme_classic(base_size = 15)+
    ggtitle(MapResults$NAME[indToPlot])
  print(ggPlot)
  
  ggBox <- ggplot(df,aes(factor(genotype),rbon_area, col = factor(genotype)))+ geom_boxplot(show.legend = FALSE) +
    scale_color_manual(values=mypalette) +
    theme_classic(base_size = 15) + xlab("genotype") +
    ggtitle(MapResults$NAME[indToPlot])
  print(ggBox)
}
```

```
# Set the SNP we plot
indSNP <- c(which(MapResults$NAME %in% "OAR6_40311379"))

# Call Fun_PlotDThPath
Fun_PlotDThPath(sortData,Nmpz,ThPaths,MapResults,indSNP)
```

```
# Call Fun_plotEffectSize
Fun_plotEffectSize(sortData,MapResults,indSNP)
```

#### Figure 7F

##### Functionality of top SNPs by GWAS and GIFT above \(99\%\) threshold

```
# Read the dataset of highly significant SNPs consisting of the genetic information
AllSNPsOfInterest <- read.csv("FunctionSNPsAboveThreshold099CHRM6.csv")
```
