## Supplementary material for "Investigative power of Genomic Informational Field Theory (GIFT) relative to GWAS for genotype-phenotype mapping": S9

Sheep Analysis: Run GIFT with residuals of model using genotype coming from filters by GWAS


### Sheep Analysis: Run GIFT with residuals of model using genotype coming from filters by GWAS

###### Supplemental material by Kyratzi et al.

```
#### CODE to run in terminal ####

# set the working directory
cd set/your/path 
  
# Load PLINK
module load plink

# Filter the dataset
plink --file texel --noweb --allow-no-sex --autosome-num 26  --recode --transpose --out texel_349
plink --noweb --tfile texel_349  --geno 0.10  --hwe 0.000001  --maf 0.05  --autosome-num 26 --allow-no-sex --make-bed --out carbon.tfull05

# Generate .ped and .map files
plink --bfile carbon.tfull05 --recode --autosome-num 26 --out carbon.tfull05
```

### Perform GWAS analysis

```
# Perform GWAS analysis

# Calculate the relationship matrix of the individuals
./gemma-0.98.5 -bfile carbon.tfull05 -maf 0 -miss 0.1 -gk 1 -o Rmatrix

# The residuals of the natural log-transformed metabolites SAM, mB12, aB12, DMG, TMG, MMA, PPA are on the 7th to 13th column of the .ped file
./gemma-0.98.5 -bfile carbon.tfull05 -n 2 -hwe 0 -maf 0 -miss 1 -k Rmatrix.cXX.txt -lmm 4 -o GWAS_rlnSAM_05
./gemma-0.98.5 -bfile carbon.tfull05 -n 3 -hwe 0 -maf 0 -miss 1 -k Rmatrix.cXX.txt -lmm 4 -o GWAS_rlnmB12_05
./gemma-0.98.5 -bfile carbon.tfull05 -n 4 -hwe 0 -maf 0 -miss 1 -k Rmatrix.cXX.txt -lmm 4 -o GWAS_rlnaB12_05
./gemma-0.98.5 -bfile carbon.tfull05 -n 5 -hwe 0 -maf 0 -miss 1 -k Rmatrix.cXX.txt -lmm 4 -o GWAS_rlnDMG_05
./gemma-0.98.5 -bfile carbon.tfull05 -n 6 -hwe 0 -maf 0 -miss 1 -k Rmatrix.cXX.txt -lmm 4 -o GWAS_rlnTMG_05
./gemma-0.98.5 -bfile carbon.tfull05 -n 7 -hwe 0 -maf 0 -miss 1 -k Rmatrix.cXX.txt -lmm 4 -o GWAS_rlnMMA_05
./gemma-0.98.5 -bfile carbon.tfull05 -n 8 -hwe 0 -maf 0 -miss 1 -k Rmatrix.cXX.txt -lmm 4 -o GWAS_rlnPPA_05

# Export GWAS results
# Export columns of chromosome, name, position and pwald
cd output
cat GWAS_rlnSAM_05.assoc.txt | awk '{print $1, $2, $3, $13}' > GWAS_rlnSAM_05.assoc_subset.txt
cat GWAS_rlnmB12_05.assoc.txt | awk '{print $1, $2, $3, $13}' > GWAS_rlnmB12_05.assoc_subset.txt
cat GWAS_rlnaB12_05.assoc.txt | awk '{print $1, $2, $3, $13}' > GWAS_rlnaB12_05.assoc_subset.txt
cat GWAS_rlnDMG_05.assoc.txt | awk '{print $1, $2, $3, $13}' > GWAS_rlnDMG_05.assoc_subset.txt
cat GWAS_rlnTMG_05.assoc.txt | awk '{print $1, $2, $3, $13}' > GWAS_rlnTMG_05.assoc_subset.txt
cat GWAS_rlnMMA_05.assoc.txt | awk '{print $1, $2, $3, $13}' > GWAS_rlnMMA_05.assoc_subset.txt
cat GWAS_rlnPPA_05.assoc.txt | awk '{print $1, $2, $3, $13}' > GWAS_rlnPPA_05.assoc_subset.txt
```

```
# Calculate the number of independent SNPs in the dataset

module load plink
plink -tfile texel_349 --noweb --allow-no-sex --autosome-num 26 --recode vcf-iid --out data
# --noweb: This flag disables web-based operations.
# --allow-no-sex: This flag allows samples with missing sex information.
# --autosome-num 26: This sets the number of autosomes in the dataset to 26. The autosomes are the non-sex chromosomes.
# --recode vcf-iid: This option specifies the output format. In this case, it's requesting the data to be recoded in the VCF (Variant Call Format) with individual identifiers.
# --out data: This specifies the prefix for the output files.

module load bcftools
bcftools +prune --max r2=0.1 --window 1Mb -Ov -o testoutput.vcf data.vcf
# +prune: This option is used to perform LD (linkage disequilibrium) pruning on the input VCF file. LD pruning is a common step in population genetics to reduce redundancy in genetic data by removing correlated variants.
# --max r2=0.1: This sets the maximum r-squared (a measure of correlation between genetic variants) allowed for variants to be considered correlated. Variants with an r-squared above 0.1 will be pruned (removed).
# --window 1Mb: This defines the size of the sliding window for LD pruning. Variants within this window will be considered for LD pruning. In this case, the window size is set to 1 megabase (1Mb).
# -Ov: This option specifies the output format as VCF.
# -o testoutput.vcf: This specifies the output file name. The pruned VCF data will be saved to a file named testoutput.vcf.
# data.vcf: This is the input VCF file on which LD pruning will be performed.

bcftools stats testoutput.vcf > testoutput.vcf.stats
# stats: The subcommand that computes various statistics for a VCF file.
# testoutput.vcf: The input VCF file for which statistics are being computed.
```

### Perform GIFT analysis

```
# set the working directory
# setwd(YourPath)

# Read the ped files
require(data.table) #for fread
carbon.tfull05 <- fread("carbon.tfull05.ped", header = FALSE)
carbon.tfull05.map <- fread("carbon.tfull05.map", header = FALSE)
```

```
# Read the pheno dataset
Residuals_asPheno <- read.csv("All_ln_residuals.csv"))

# match the individuals in the the two datasets
Residuals_asPheno <- Residuals_asPheno[match(unlist(carbon.tfull05[,2]),Residuals_asPheno$id),]
```

```
# transfer the combination of alleles to microstates -1, +1, 0
# GG=CC=-1
# TT=AA=+1
# AG=CG=AC=AT=0

library(pegas)
genotype <- alleles2loci(carbon.tfull05[,-c(1:6)])

library(dplyr)
Geno <- as.data.frame(matrix(case_when((genotype=="G/G") | (genotype=="C/C") ~ -1,
                                       (genotype=="T/T") | (genotype=="A/A") ~ +1,
                                       genotype=="0/0" ~ NA_real_,
                                       TRUE ~ 0),
                             nrow = nrow(genotype), ncol = ncol(genotype), byrow = FALSE))

# Set the row and column names of the new geno dataset
rownames(Geno) <- unlist(carbon.tfull05[,2])
colnames(Geno) <- unlist(carbon.tfull05.map[,2])
```

#### Treat missing phenotypic values

```
# Function to remove the individuals with missing values on the phenotype (residuluals_ln) from phenotype list and genotype dataset
# Input variables:
#   Phenotype: dataframe consisting of info about individuals and the values of metabolites
#   Genotype: dataframe consisting of microstates (row: individuals, col: SNPs)
# Output variables:
#   List of phenotype and genotype datasets (row: individuals, col: phenotype) after removing individuals with missing phenotype

Fun_filterMissingPheno <- function(Phenotype, Genotype, namePheno){
  ind.missPheno <- which(is.na(Phenotype[,namePheno]))
  Phenotype <- Phenotype[-ind.missPheno,c("id", namePheno)]
  Genotype <- Genotype[-ind.missPheno,]
  
  return(list(Pheno = Phenotype, Geno = Genotype))
}

# Call Fun_filterMissingPheno
# Data_noMissPheno <- Fun_filterMissingPheno(Residuals_asPheno, Geno, "rlnSam")
# Data_noMissPheno <- Fun_filterMissingPheno(Residuals_asPheno, Geno, "rlnmB12")
# Data_noMissPheno <- Fun_filterMissingPheno(Residuals_asPheno, Geno, "rlnaB12")
# Data_noMissPheno <- Fun_filterMissingPheno(Residuals_asPheno, Geno, "rlnDMG")
# Data_noMissPheno <- Fun_filterMissingPheno(Residuals_asPheno, Geno, "rlnTMG")
# Data_noMissPheno <- Fun_filterMissingPheno(Residuals_asPheno, Geno, "rlnMMA")
# Data_noMissPheno <- Fun_filterMissingPheno(Residuals_asPheno, Geno, "rlnPPA")
```

```
# Function to compute the cumulative distributions of the ordered and random configuration according to the bins for each SNP
# Input variables:
#   Geno: sorted microstate dataset (row: individuals, col: SNPs)
#   CountValBins:  dataframe with the number of how many individuals in each bin (rows) per SNP (columns)
#   Nmpz: dataframe of number of microstates for each SNP (rows)
# Output variables:
#   List of Wp, Wm, Wz, W0p, W0m, W0z: Cumulative distributions of +1,-1 respectively for ordered and random configuration according to the bins for each SNP

Fun_CumDistr <- function(Geno, CountValBins, Nmpz){
  library(purrr) # for accumulate
  
  CumDistrW <- Map(function(Geno,CountValBins){
    bins <- CountValBins[!is.na(CountValBins)]
    numBins <- length(bins)
    
    # Split the vector of microstates into groups per bin
    grouped_geno <- split(Geno[!is.na(Geno)], rep(seq_along(bins), bins))
    # Remove NA values from each vector in the list
    grouped_geno <- lapply(grouped_geno, function(x) na.omit(x))
    
    # Define a function to calculate the weighted cumulative sums
    calculate_weighted_cumulative_sums_per_bin <- function(grouped_geno) {
      weights <- (1:length(grouped_geno)) / length(grouped_geno)
      group_cumsum <- sapply(-1:1, function(group) {
        weights * sum(grouped_geno == group)
      })
      return(group_cumsum)
    }
    
    # Calculate the weighted cumulative sums for each group per bin
    cumulative_sums <- t(matrix(sapply(grouped_geno, calculate_weighted_cumulative_sums_per_bin)))
    cumulative_sums <- do.call(rbind, lapply(cumulative_sums, function(x){matrix(x, ncol = 3, byrow = FALSE)}))
    
    # Add the cumulative sums from previous bins
    result <- apply(cumulative_sums, 2, function(x){
      accumulate(2:numBins, function(x, n) {
        indstart <- sum(bins[1:(n-1)]) + 1
        indend <- sum(bins[1:n])
        s <- x[indstart - 1]
        x[indstart:indend] <- x[indstart:indend] + s
        return(x)}, .init = x)})
    
    Wm <- result[[1]][[length(result[[1]])]]
    Wz <- result[[2]][[length(result[[2]])]]
    Wp <- result[[3]][[length(result[[3]])]]
    
    return(data.frame(Wp = Wp, Wm = Wm, Wz = Wz))
    
  }, Geno, CountValBins)
  
  Wp <- as.data.frame(do.call(qpcR:::cbind.na, lapply(CumDistrW, `[[`, "Wp")))
  Wm <- as.data.frame(do.call(qpcR:::cbind.na, lapply(CumDistrW, `[[`, "Wm")))
  Wz <- as.data.frame(do.call(qpcR:::cbind.na, lapply(CumDistrW, `[[`, "Wz")))
  
  CumDistrW0 <- apply(Nmpz, 1, function(Nmpz){
    Nmpz <- unlist(Nmpz)
    N <- sum(Nmpz)
    W0p <- c(1:N) * Nmpz[2] / N
    W0m <- c(1:N) * Nmpz[1] / N
    W0z <- c(1:N) * Nmpz[3] / N
    return(data.frame(W0p = W0p, W0m = W0m, W0z = W0z))
  })
  
  W0p <- as.data.frame(do.call(qpcR:::cbind.na, lapply(CumDistrW0, `[[`, "W0p")))
  W0m <- as.data.frame(do.call(qpcR:::cbind.na, lapply(CumDistrW0, `[[`, "W0m")))
  W0z <- as.data.frame(do.call(qpcR:::cbind.na, lapply(CumDistrW0, `[[`, "W0z")))
  
  return(list(Wp = Wp, Wm = Wm, Wz = Wz, W0p = W0p, W0m = W0m, W0z = W0z))
}


# Call Fun_CumDistr
CumDistr <- Fun_CumDistr(sortData$Geno, CountValBins, Nmpz)
```

```
# Function to calculate Theta-paths
# Input variables:
#   CumDistr: List consisting of the Cumulative Distributions Wp, Wm, Wz, W0p, W0m, W0z
# Output variables:
#   List of Thp = Wp - W0p,
#           Thm = Wm - W0m,
#           Thj = Wp - Wm (the phenotype-responding genetic path),
#           Th0j = W0p - W0m (the default genetic path),
#           DThj = Thj - Th0j

Fun_ThPaths <- function(CumDistr){
  
  N <- nrow(CumDistr$Wp)
  num_SNPs <- ncol(CumDistr$Wp)
  
  Thp <- data.frame(matrix(NA, nrow = N, ncol = num_SNPs))
  Thm <- data.frame(matrix(NA, nrow = N, ncol = num_SNPs))
  Thj <- data.frame(matrix(NA, nrow = N, ncol = num_SNPs))
  Th0j <- data.frame(matrix(NA, nrow = N, ncol = num_SNPs))
  DThj <- data.frame(matrix(NA, nrow = N, ncol = num_SNPs))
  
  Thp <- CumDistr$Wp - CumDistr$W0p
  Thm <- CumDistr$Wm -CumDistr$W0m
  Thj <- CumDistr$Wp - CumDistr$Wm
  Th0j <- CumDistr$W0p - CumDistr$W0m  
  DThj <- Thj - Th0j
  
  return(list(Thp = Thp, Thm = Thm, Thj = Thj, Th0j = Th0j, DThj = DThj))
}


# Call Fun_CumDistr
ThPaths <- Fun_ThPaths(CumDistr)
```

#### pGIFT for two and three states

For +1,-1:

\[pGIFT =
\Bigg(\frac{1}{2}-\frac{1}{\pi}tan^{-1} \Big(\frac{\sqrt{-2\phi
\tilde\phi}}{2(\phi-\tilde\phi)} \Big) \Bigg) \exp\Big\{
-\frac{(\phi-\tilde\phi)^2}{2Nw(1-w)} \Big\}, w=N\_+/N, \quad \phi =
\max\_{j}\Delta\Theta(j) , \quad \tilde\phi = \min\_{j}\Delta\Theta(j)
\]

\[=
\Bigg(\frac{1}{2}-\frac{1}{\pi}tan^{-1} \Big(\frac{\sqrt{-2\phi
\tilde\phi}}{2(\phi-\tilde\phi)} \Big) \Bigg) \exp\Big\{
-\frac{N(\phi-\tilde\phi)^2}{2N\_+N\_-} \Big\} \]

For +1,0:

\[pGIFT =
\Bigg(\frac{1}{2}-\frac{1}{\pi}tan^{-1} \Big(\frac{\sqrt{-2\phi
\tilde\phi}}{2(\phi-\tilde\phi)} \Big) \Bigg) \exp\Big\{
-\frac{2(\phi-\tilde\phi)^2}{Nw(1-w)} \Big\}, w=N\_+/N, \quad \phi =
\max\_{j}\Delta\Theta(j) , \quad \tilde\phi = \min\_{j}\Delta\Theta(j)
\]

\[=
\Bigg(\frac{1}{2}-\frac{1}{\pi}tan^{-1} \Big(\frac{\sqrt{-2\phi
\tilde\phi}}{2(\phi-\tilde\phi)} \Big) \Bigg) \exp\Big\{
-\frac{2N(\phi-\tilde\phi)^2}{N\_+N\_0} \Big\} \]

For -1,0:

\[pGIFT =
\Bigg(\frac{1}{2}-\frac{1}{\pi}tan^{-1} \Big(\frac{\sqrt{-2\phi
\tilde\phi}}{2(\phi-\tilde\phi)} \Big) \Bigg) \exp\Big\{
-\frac{2(\phi-\tilde\phi)^2}{Nw(1-w)} \Big\}, w=N\_-/N, \quad \phi =
\max\_{j}\Delta\Theta(j) , \quad \tilde\phi =
\min\_{j}\Delta\Theta(j)\] \[ =
\Bigg(\frac{1}{2}-\frac{1}{\pi}tan^{-1} \Big(\frac{\sqrt{-2\phi
\tilde\phi}}{2(\phi-\tilde\phi)} \Big) \Bigg) \exp\Big\{
-\frac{2N(\phi-\tilde\phi)^2}{N\_-N\_0} \Big\}\]

For -1,+1,0:

\[pGIFT =
\Bigg(\frac{1}{2}-\frac{1}{\pi}tan^{-1} \Big(\frac{\sqrt{-\phi
\tilde\phi}}{(\phi-\tilde\phi) \sqrt{2}} \Big) \Bigg) \exp\Big\{
-\frac{8N(\phi-\tilde\phi)^2}{N(N-N\_0)-(N\_+-N\_-)^2} \Big\}, \quad \phi =
\max\_{j}\Delta\Theta(j) , \quad \tilde\phi =
\min\_{j}\Delta\Theta(j)\]

```
# Function to calculate the p-value
# Input variables:
#   Geno: sorted geno dataset (row: individuals, col: SNPs)
#   Nmpz: dataframe of number of microstates for each SNP
#   ThPaths: List of Theta-paths (output of Fun_ThPaths)
# Output variables:
#   pGIFT: vector consists of pGIFT values

Fun_pGIFT <- function(Geno, Nmpz, ThPaths){
  pGIFT <- c()
  
  num_SNPs <- ncol(Geno)
  DThj <- as.data.frame(ThPaths$DThj)
  
  phi <- apply(DThj, 2, function(x){max(x, na.rm = TRUE)})
  phi_tilde <- apply(DThj, 2, function(x){min(x, na.rm = TRUE)})
  
  for (i in 1:num_SNPs) {
    
    no.NA <- !is.na(Geno[,i])
    N <- sum(no.NA)
    
    if (sum(Nmpz[i,]>=3)==3) { #check if we have three states - compute the pGIFT for three-states
      e <- exp( -(8*N*(phi[i]-phi_tilde[i])^2) / (N*(N-Nmpz[i,3])-(Nmpz[i,2]-Nmpz[i,1])^2) )
      
      pGIFT[i] <- ( (1/2)-(1/pi)*atan(sqrt(-phi[i]*phi_tilde[i])/((phi[i]-phi_tilde[i])*sqrt(2))) ) *e
      
    } #end if condition for three states
    if (sum(Nmpz[i,]>=3)==2) {  # length(uniq.states)==2
      uniq.states <- data.frame(table(Geno[no.NA,i]))
      ind_geno <- which(uniq.states$Freq>=3)
      if (length(intersect(c(-1,0),uniq.states[ind_geno,1]))==2){ #check if we have -1 and 0 states to use Thjm and Thjz
        e <- exp( -(2*N*(phi[i]-phi_tilde[i])^2) / (uniq.states$Freq[ind_geno[1]]*uniq.states$Freq[ind_geno[2]]) )
        
        pGIFT[i] <- ( (1/2)-(1/pi)*atan(sqrt(-2*phi[i]*phi_tilde[i])/(2*(phi[i]-phi_tilde[i]))) ) *e
      } #end if condition for two states: -1, 0
      
      if (length(intersect(c(+1,0),uniq.states[ind_geno,1]))==2){ #check if we have +1 and 0 states to use Thjp and Thjz
        e <- exp( -(2*N*(phi[i]-phi_tilde[i])^2) / (uniq.states$Freq[ind_geno[1]]*uniq.states$Freq[ind_geno[2]]) )
        
        pGIFT[i] <- ( (1/2)-(1/pi)*atan(sqrt(-2*phi[i]*phi_tilde[i])/(2*(phi[i]-phi_tilde[i]))) ) *e
      } #end if condition for two states: +1, 0
      
      else if (length(intersect(c(-1,+1),uniq.states[ind_geno,1]))==2){ #check if we have -1 and +1 states to use Thjm and Thjp
        e <- exp( -(N*(phi[i]-phi_tilde[i])^2) / (2*uniq.states$Freq[ind_geno[1]]*uniq.states$Freq[ind_geno[2]]) )
        
        pGIFT[i] <- ( (1/2)-(1/pi)*atan(sqrt(-2*phi[i]*phi_tilde[i])/(2*(phi[i]-phi_tilde[i]))) ) *e
      } #end if condition for two states: -1, +1
    } #end if condition for two states
    
    if (sum(Nmpz[i,]>=3)<2) { #check if we have only one state
      pGIFT[i] <- 1 # set this to one so after taking the logarithm we have zero (non-significant)
    }
    
  }
  
  return(pGIFT)
  
}

# Call Fun_pGIFT
pGIFT <- Fun_pGIFT(sortData$Geno, Nmpz, ThPaths)

# Adjust pGIFT based on Benjamini Hochberg procedure
pGIFT_adjBH <- p.adjust(pGIFT, method = "BH")
```

### RESULTS

#### Figure 8A

```
# Read GWAS results
pGWAS_rlnSAM <- read.table("GWAS_rsam_05.assoc_subset.txt", header = TRUE)
pGWAS_rlnSAM$mlog10_pGWAS <- -log10(pGWAS_rlnSAM$p_wald)

pGWAS_rlnmB12 <- read.table("GWAS_rmB12_05.assoc_subset.txt", header = TRUE)
pGWAS_rlnmB12$mlog10_pGWAS <- -log10(pGWAS_rlnmB12$p_wald)

pGWAS_rlnaB12 <- read.table("GWAS_raB12_05.assoc_subset.txt", header = TRUE)
pGWAS_rlnaB12$mlog10_pGWAS <- -log10(pGWAS_rlnaB12$p_wald)

pGWAS_rlnDMG <- read.table("GWAS_rDMG_05.assoc_subset.txt", header = TRUE)
pGWAS_rlnDMG$mlog10_pGWAS <- -log10(pGWAS_rlnDMG$p_wald)

pGWAS_rlnTMG <- read.table("GWAS_rTMG_05.assoc_subset.txt", header = TRUE)
pGWAS_rlnTMG$mlog10_pGWAS <- -log10(pGWAS_rlnTMG$p_wald)

pGWAS_rlnMMA <- read.table("GWAS_rMMA_05.assoc_subset.txt", header = TRUE)
pGWAS_rlnMMA$mlog10_pGWAS <- -log10(pGWAS_rlnMMA$p_wald)

pGWAS_rlnPPA <- read.table("GWAS_rPPA_05.assoc_subset.txt", header = TRUE)
pGWAS_rlnPPA$mlog10_pGWAS <- -log10(pGWAS_rlnPPA$p_wald)
```

```
# Read GIFT results
require(data.table)
MapResults_rlnSAM <- fread("MapResults_rlnSAM.csv")
MapResults_rlnmB12 <- fread("MapResults_rlnmB12.csv")
MapResults_rlnaB12 <- fread("MapResults_rlnaB12.csv")
MapResults_rlnDMG <- fread("MapResults_rlnDMG.csv")
MapResults_rlnTMG <- fread("MapResults_rlnTMG.csv")
MapResults_rlnMMA <- fread("MapResults_rlnMMA.csv")
MapResults_rlnPPA <- fread("MapResults_rlnPPA.csv")
```

```
# Read the thresholds values as calculated by the permutation analysis
threshold099 <-readRDS("threshold099N565.rds")
threshold095 <-readRDS("threshold095N565.rds")
```

```
library(qqman)
# Function to generate Manhattan plot for GWAS analyses
Fun_GWASManhattanPlots <- function(pGWAS_rln, metabolite){
  num_IndepSNPs <- 624 # set the number of independent SNPs as calculated using bcftools
  manhattan(pGWAS_rln, chr = "chr", bp = "ps", p = "mlog10_pGWAS", snp = "rs",
            chrlabs = as.character(unique(pGWAS_rln$chr)), logp = FALSE, ylab = "-log10(pGWAS)", 
            cex = 0.8, cex.axis = 2, cex.lab = 2, cex.main = 2, ylim = c(0,10),  col = c("blue3", "orange3"),
            main = paste0("GWAS_rln", metabolite), suggestiveline = FALSE, genomewideline = FALSE)
  abline(h = -log10(0.01/num_IndepSNPs), col = "red", lty = "dashed", lwd = 2)
  abline(h = -log10(0.05/num_IndepSNPs), col = "red", lty = "dashed", lwd = 2)
}

# Function to generate Manhattan plot for GIFT analyses
Fun_GIFTManhattanPlots <- function(MapResults_rln, metabolite, threshold1, threshold2){
manhattan(MapResults_rln, chr = "CHRM", bp = "POSITION", p = "mlog10_pGIFTadjBH", snp = "NAME",
            chrlabs = as.character(unique(MapResults_rln$CHRM)), logp = FALSE, ylab = "-log10(pGIFT)", 
          cex = 0.8, cex.axis = 2, cex.lab = 2, cex.main = 2, ylim = c(0,25),  col = c("blue3", "orange3"),
            main = paste0("GIFT_rln", metabolite), suggestiveline = FALSE, genomewideline = FALSE)
  abline(h = threshold1, col = "red", lty = "dashed", lwd = 2)
  abline(h = threshold2, col = "red", lty = "dashed", lwd = 2)
}

# Call Fun_GWASManhattanPlots and Fun_GIFTManhattanPlots
par(mfrow=c(7,2))

Fun_GWASManhattanPlots(pGWAS_rlnSAM, "SAM")
Fun_GIFTManhattanPlots(MapResults_rlnSAM, "SAM", threshold099, threshold095)

Fun_GWASManhattanPlots(pGWAS_rlnmB12, "mB12")
Fun_GIFTManhattanPlots(MapResults_rlnmB12, "mB12", threshold099, threshold095)

Fun_GWASManhattanPlots(pGWAS_rlnaB12, "aB12")
Fun_GIFTManhattanPlots(MapResults_rlnaB12, "aB12", threshold099, threshold095)

Fun_GWASManhattanPlots(pGWAS_rlnDMG, "DMG")
Fun_GIFTManhattanPlots(MapResults_rlnDMG, "DMG", threshold099, threshold095)

Fun_GWASManhattanPlots(pGWAS_rlnTMG, "TMG")
Fun_GIFTManhattanPlots(MapResults_rlnTMG, "TMG", threshold099, threshold095)

Fun_GWASManhattanPlots(pGWAS_rlnMMA, "MMA")
Fun_GIFTManhattanPlots(MapResults_rlnMMA, "MMA", threshold099, threshold095)

Fun_GWASManhattanPlots(pGWAS_rlnPPA, "PPA")
Fun_GIFTManhattanPlots(MapResults_rlnPPA, "PPA", threshold099, threshold095)
```

#### Figure 8B

##### Functionality of top SNPs by GIFT above \(99\%\) threshold

```
# Read the dataset of highly significant SNPs consisting of the genetic information
GIFThighSNPs_SAM <- read.csv("GIFThighSNPs_SAM.csv")
GIFThighSNPs_mB12 <- read.csv("GIFThighSNPs_mB12.csv")
GIFThighSNPs_aB12 <- read.csv("GIFThighSNPs_aB12.csv")
GIFThighSNPs_DMG <- read.csv("GIFThighSNPs_DMG.csv")
GIFThighSNPs_TMG <- read.csv("GIFThighSNPs_TMG.csv")
GIFThighSNPs_MMA <- read.csv("GIFThighSNPs_MMA.csv")
GIFThighSNPs_PPA <- read.csv("GIFThighSNPs_PPA.csv")
```

```
# Generate the barplots
library(ggplot2)

## SAM
ggSAM <- ggplot(GIFThighSNPs_SAM, aes(x = ConsequenceType)) +
  geom_bar() +
  xlab("Consequence Type") +
  ylab("SNPs per Consequence Type") +
  ylim(0,15) +
  theme_bw() +
  ggtitle("GIFT_SAM") +
  theme(text = element_text(size = 20),
        axis.text = element_text(color = "black"),
        title = element_text(size = 20),
        axis.text.x = element_text(angle = 35, vjust = 1, hjust=1)) +
  scale_x_discrete(labels=c('3 prime UTR', 'intron','missense', 'synonymous'))

## mB12
ggmB12 <- ggplot(GIFThighSNPs_mB12, aes(x = ConsequenceType)) +
  geom_bar() +
  xlab("Consequence Type") +
  ylab("SNPs per Consequence Type") +
  ylim(0,15) +
  theme_bw() +
  ggtitle("GIFT_mB12") +
  theme(text = element_text(size = 20),
        axis.text = element_text(color = "black"),
        title = element_text(size = 20),
        axis.text.x = element_text(angle = 35, vjust = 1, hjust=1)) +
  scale_x_discrete(labels=c('missense', 'synonymous', 'upstream'))

## aB12
ggaB12 <- ggplot(GIFThighSNPs_aB12, aes(x = ConsequenceType)) +
  geom_bar() +
  xlab("Consequence Type") +
  ylab("SNPs per Consequence Type") +
  ylim(0,15) +
  theme_bw() +
  ggtitle("GIFT_aB12") +
  theme(text = element_text(size = 20),
        axis.text = element_text(color = "black"),
        title = element_text(size = 20),
        axis.text.x = element_text(angle = 35, vjust = 1, hjust=1)) +
  scale_x_discrete(labels=c('intron','synonymous', 'upstream', 'NA'))

## DMG
ggDMG <- ggplot(GIFThighSNPs_DMG, aes(x = ConsequenceType)) +
  geom_bar() +
  xlab("Consequence Type") +
  ylab("SNPs per Consequence Type") +
  ylim(0,30) +
  theme_bw() +
  ggtitle("GIFT_DMG") +
  theme(text = element_text(size = 20),
        axis.text = element_text(color = "black"),
        title = element_text(size = 20),
        axis.text.x = element_text(angle = 35, vjust = 1, hjust=1)) +
  scale_x_discrete(labels=c('3 prime UTR', 'intron','missense',  'splice donor region', 
                            'splice region', 'synonymous', 'upstream', 'NA'))

## TMG
ggTMG <- ggplot(GIFThighSNPs_TMG, aes(x = ConsequenceType)) +
  geom_bar() +
  xlab("Consequence Type") +
  ylab("SNPs per Consequence Type") +
  ylim(0,20) +
  theme_bw() +
  ggtitle("GIFT_TMG") +
  theme(text = element_text(size = 20),
        axis.text = element_text(color = "black"),
        title = element_text(size = 20),
        axis.text.x = element_text(angle = 35, vjust = 1, hjust=1)) +
  scale_x_discrete(labels=c('intron','missense','synonymous', 'upstream', 'NA'))

## MMA
ggMMA <- ggplot(GIFThighSNPs_MMA, aes(x = ConsequenceType)) +
  geom_bar() +
  xlab("Consequence Type") +
  ylab("SNPs per Consequence Type") +
  ylim(0,8) +
  theme_bw() +
  ggtitle("GIFT_MMA") +
  theme(text = element_text(size = 20),
        axis.text = element_text(color = "black"),
        title = element_text(size = 20),
        axis.text.x = element_text(angle = 35, vjust = 1, hjust=1)) +
  scale_x_discrete(labels=c('3 prime UTR', 'intron', 'synonymous', 'upstream'))

## PPA
ggPPA <- ggplot(GIFThighSNPs_PPA, aes(x = ConsequenceType)) +
  geom_bar() +
  xlab("Consequence Type") +
  ylab("SNPs per Consequence Type") +
  ylim(0,40) +
  theme_bw() +
  ggtitle("GIFT_PPA") +
  theme(text = element_text(size = 20),
        axis.text = element_text(color = "black"),
        title = element_text(size = 20),
        axis.text.x = element_text(angle = 35, vjust = 1, hjust=1)) +
  scale_x_discrete(labels=c('3 prime UTR', 'intron','missense', 'splice donor 5th base', 
                            'synonymous', 'upstream', 'NA'))

library(gridExtra)
grid.arrange(grobs = list(ggSAM, ggmB12, ggaB12, ggDMG, ggTMG, ggMMA, ggPPA), ncol = 2)
```
